# Plasma glycoproteomics delivers high-specificity disease biomarkers by detecting site-specific glycosylation abnormalities

**DOI:** 10.1101/2022.05.31.494121

**Authors:** Hans JCT Wessels, Purva Kulkarni, Maurice van Dael, Anouk Suppers, Esther Willems, Fokje Zijlstra, Else Kragt, Jolein Gloerich, Pierre-Olivier Schmit, Stuart Pengelley, Kristina Marx, Alain J van Gool, Dirk J Lefeber

## Abstract

The human plasma glycoproteome holds enormous potential to identify personalized biomarkers to diagnose and understand disease. Recent advances in mass spectrometry and software development are opening novel avenues to mine the glycoproteome for protein- and site-specific glycosylation changes. Here, we describe a novel plasma N-glycoproteomics method for disease diagnosis and evaluated its clinical applicability by performing comparative glycoproteomics in blood plasma of 40 controls and a cohort of 74 patients with 13 different genetic diseases that directly impact the protein N-glycosylation pathway. The plasma glycoproteome yielded high-specificity biomarker signatures for each of the individual genetic defects. Bioinformatic analyses revealed site-specific glycosylation differences that could be explained by underlying glycobiology and in specific diseases by protein-intrinsic factors. Our work illustrates the strong potential of plasma glycoproteomics to significantly increase specificity of glycoprotein biomarkers with direct insights in site-specific glycosylation changes to better understand the mechanisms underlying human disease.

## Introduction

Protein glycosylation is one of the most prominent post-translational modifications known, with strong effects on protein biology^1^. The blood plasma glycoproteome holds great potential for biomarker discovery since abnormal glycomes have been reported for numerous human diseases^2^. Evidence is emerging that protein-specific analysis of glycosylation changes will drastically increase the specificity of such biomarkers^3–6^. Proteome-wide analysis of protein glycosylation through liquid chromatography – tandem mass spectrometry (LC-MS/MS) analysis of glycopeptides, or glycoproteomics, has matured to such a state in recent years that clinical application is becoming reality to improve diagnostics and gain new insights into disease mechanisms.

Recent efforts have been made to optimize glycoproteomics technology for blood plasma samples using different tandem mass spectrometry approaches for N-glycopeptide and O-glycopeptide analysis^7–9^. Significant advances in hardware and software developments have been made, while methods for correct elucidation of both glycan and peptide moieties from large numbers of MS/MS fragmentation spectra are emerging^10^. This pioneering work yielded a characterization depth of the N- and O-glycoproteome in plasma ranging from tens of glycosylation sites and glycoproteins up to hundreds of glycosylation sites and glycoproteins depending on the sample fractionation depth and thus measurement time per sample^11–17^. As such, individual glycopeptide differentials have started to be identified in diseases such as cancer^18–22^, bacterial bloodstream infection^23^, IgA nephropathy^24^ and myocardial infarction^25^. These studies focused on glycopeptides as individual biomarkers by direct comparison of their signal intensities between sample groups which can be influenced by e.g. changes in expression level of the protein carrier, glycoform shifts or even additional post-translational modifications of the peptide-moiety. This leaves the key question if plasma glycopeptide differentials are clinically relevant to enable interpretation of underlying glycobiology in disease.

To answer this question we developed a glycoproteomics method based on glycopeptide profiling in blood plasma and assessed its clinical applicability for disease diagnosis by investigating a large cohort of patients with well-defined congenital disorder of glycosylation (CDG). CDG plasma samples present unique possibilities as model system to define aberrant glycosylation in the context of the underlying glycobiology due to their primary defects in N-glycan synthesis. This allowed us to demonstrate superior sample classification using glycopeptide intensities as compared to the use of single protein MS in current diagnostics. Moreover, we were able to deduce protein- and site-specific glycosylation shifts to translate to biological and clinical insights.

## Results

### Towards a clinically applicable strategy for plasma glycoproteomics

In view of the intended clinical application, we developed a holistic glycopeptide profiling workflow that required practical sample amounts (10μl of blood plasma) whilst achieving reasonable sample throughput for analysis (20 samples in 24 hours). Intact tryptic glycopeptides were enriched using Sepharose CL-4B^26^ in 96-well plates and analysed by C18 reversed phase LC-MS/MS without any further fractionation (figure 1a). Fragmentation experiments were performed at “low” and “high” collision energies to generate structurally informative fragments for the glycan- and peptide-moiety, respectively^27^. Optimal collision energy settings were selected using glycan- and peptide-database search results based on the number of identifications and identification scores. For pre-processing of acquired data we used available software to extract LC-MS feature information (OpenMS^28^) or to identify glycan compositions (ProteinScape™-GlycoQuest^29^) and peptide sequences (MASCOT^30^). MATLAB scripts were developed to integrate the output of individual software tools by mapping peptide and glycan identities onto their respective LC-MS features in the consensus map. This strategy supported the use of all quantified LC-MS features in subsequent statistical and chemometric analyses irrespective of identification status and maximized feature annotations by integrating identification results from all analyses.

**Figure 1.**
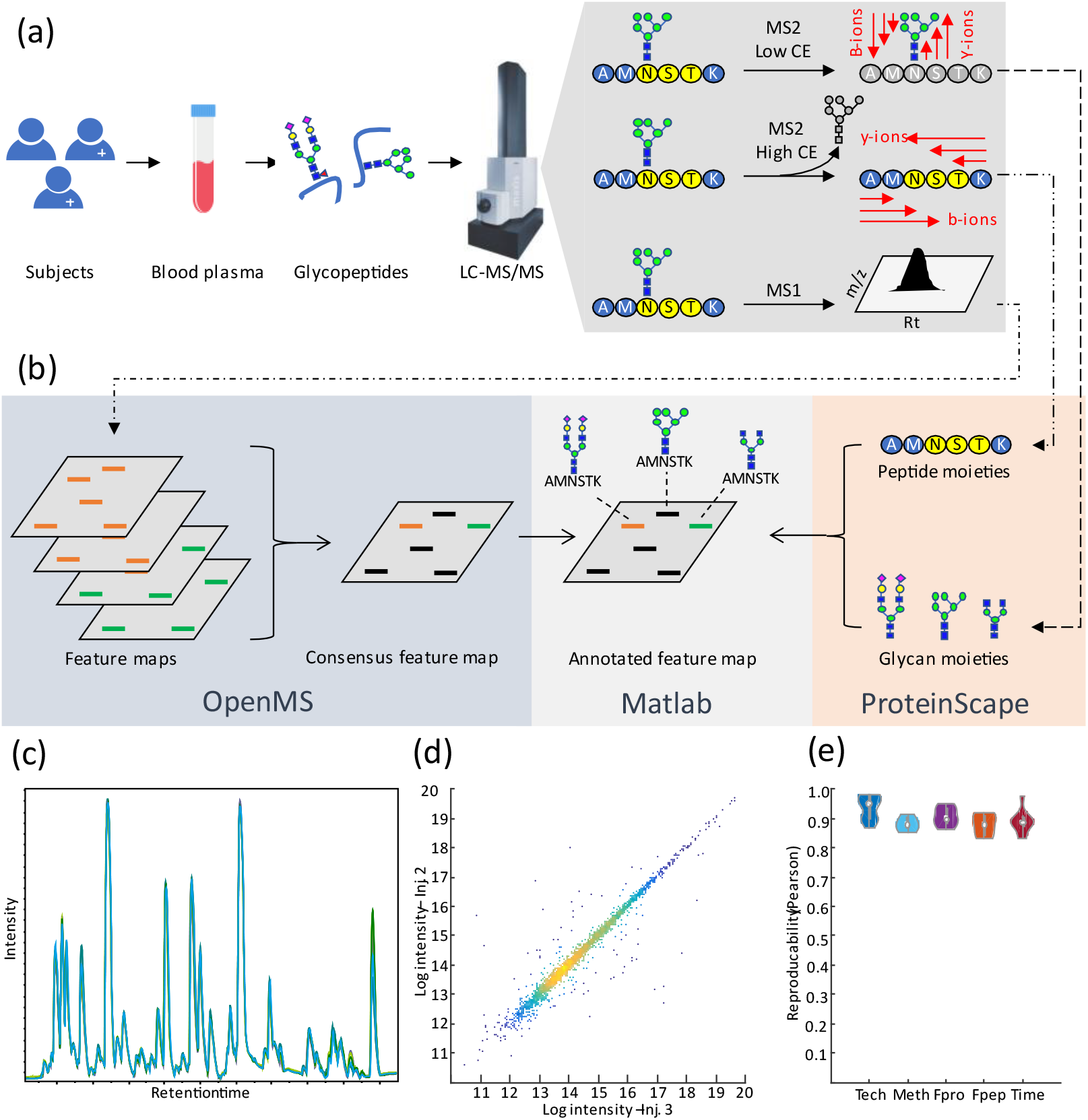
Overall glycoproteomics strategy and analytical performance. **(a)** Data generation workflow: Plasma proteins were subjected to in solution tryptic digestion and glycopeptides were enriched by solid phase extraction using Sepharose CL-4B material^26^. Glycopeptide mixtures were analysed by C18 reversed phase liquid chromatography with online tandem mass spectrometry using low and high collision energies (CE) for glycan- and peptide-moiety fragmentation experiments, respectively^27^. **(b)** Data processing workflow. Quantitative information was extracted from data files as feature maps and subsequently combined into a consensus feature map in OpenMS software. ProteinScape™ 3.1 was used to perform MS/MS glycopeptide spectrum searches and classified MS/MS spectra were searched against the CarbBank glycan database using GlycoQuest or against the Swiss-prot human protein sequence database using MASCOT. In-house developed MATLAB scripts were used to map identified glycan- and peptide-moieties onto the consensus feature map for subsequent analyses. **(c)** Base peak chromatogram overlay from five replicate injections of a glycopeptide preparation from a single control sample. **(d)** Log intensity scatterplot of features for two replicate injections of a single control sample. **(e)** Violin plot of Pearson’s correlation coefficients from replicate injections (Tech; n=5), independent sample preparations and measurements of a single biological sample (Meth; n=5), 1-5 plasma feeze/thaw cycles (Fpro n=5), 1-5 glycopeptide sample freeze/thaw cycles (Fpep n=5), and in-time 0-24 hr technical replicates of single glycopeptide samples stored at 10°C (Time; n=5).

Robust analytical performance and sample stability are prerequisites for successful clinical application of plasma glycoproteomics technology. A single control plasma sample was used to determine the intra- and inter-essay reproducibility *via* five replicate injections of the same sample preparation and analysis of five independent glycopeptide preparations of the same sample. Pooled Pearson’s correlation (PPC) values and median CV values for feature intensities were used to assess analytical performance. The feature intensities were not subjected to any signal intensity normalisation procedure to assess intrinsic intensity variability. The PPC value of 0.94 (figure 1e) and median CV of 11% for replicate injections show that analytical variability is well controlled on our LC-MS platform as can be observed from the base peak chromatogram overlays (figure 1c) and intensity scatterplot of two replicate injections (figure 1d). In inter-assay comparisons, the PPC value was 0.88 and median CV was 23%. To evaluate the effects of sample stability on the glycoproteomics workflow, we subjected a control pre- and post-digested plasma sample for up to five freeze/thaw cycles and incubated a control sample digest in the autosampler at 10°C for up to 24 hours. Results for the plasma sample freeze/thaw cycles did show a mild increase in variability based on median CV values (29%) but did not affect the correlation between samples (PPC=0.90). Incubation of a control sample in the autosampler did not increase the variability based on the PPC (0.90) and median CV(21%) values compared to inter-assay variability. Similar results were obtained for up to five freeze/thaw cycles of a single glycopeptide preparation for which the median CV (11%) was identical to replicate injections of a single sample with PPC of 0.88. Retention time reproducibility was < 0.5% CV and average mass accuracy was < 2 ppm. We conclude that the analytical performance of the glycoproteomics procedure is sufficient for application in clinical cohort studies.

### Glycopeptide identification

Plasma glycopeptide identification by LC-MS/MS is known to be challenging due to low electrospray ionization efficiency, glycoform signal dilution and poor fragmentation of the peptide moiety in collision induced dissociation experiments^27^. We implemented acetonitrile-enriched nitrogen source gas^31^ to significantly increase ionization efficiency of intact glycopeptides as compared to other organic solvents that we tested (supplementary figure 1), thereby shifting the distribution of precursor ions towards higher charge states for enhanced peptide-moiety fragmentation. Application of low collision energy conditions produced rich glycan fragmentation spectra for subsequent glycan database searches. The relatively low intensity of peptide moiety b- and y-fragment ions in MS/MS spectra under high collision energy conditions complicated protein database searches. We tackled this by MS/MS spectrum pre-processing (supplementary figure 2) by removing most of the glycan B-, Y- and internal fragment ions prior to database searching. As a result, MASCOT identification scores significantly improved by 25% and raised the total number of peptide-spectrum matches by 42% to yield 57% more identified unique peptide-moieties (supplementary figure 2).

To maximize the number of annotated glycopeptide features in the consensus feature map, we then applied the plasma glycoproteomics method to a cohort of 200 individuals that consisted of 40 healthy donors, 12 patients with normal transferrin glycosylation, 30 patients with unknown genetic defect and abnormal transferrin glycosylation and 118 patients spanning 35 genetically resolved congenital disorders of glycosylation (CDG). The broad variation in glycan structures in these CDG patients was key to identify disease-specific or otherwise low-abundant glycoforms in control samples with the aid of increased signals from accumulated glycoforms. In total, 267.394 glycan-spectrum matches were obtained through GlycoQuest searches (191 unique glycan compositions) that could be mapped onto 7.229 features of the LC-MS consensus feature map which contained merged features from all individual samples^32^. The qualitative glycome representation of detected glycan species, agglomerated in glycan traits, is shown in figure 2b. For peptide moieties, 5.988 peptide-spectrum matches were obtained for the full CDG cohort dataset which could be mapped onto 1.430 LC-MS features in the consensus feature map. Peptide sequences that resulted from missed tryptic cleavages or that lacked a predicted or known N-glycosylation sequon were removed from the dataset to yield a total of 58 unique peptide sequences identified from 34 proteins for which at least one glycoform was identified. The identified proteins belong to abundant glycoproteins in blood plasma, spanning three orders of magnitude in abundance (figure 2a).

**Figure 2.**
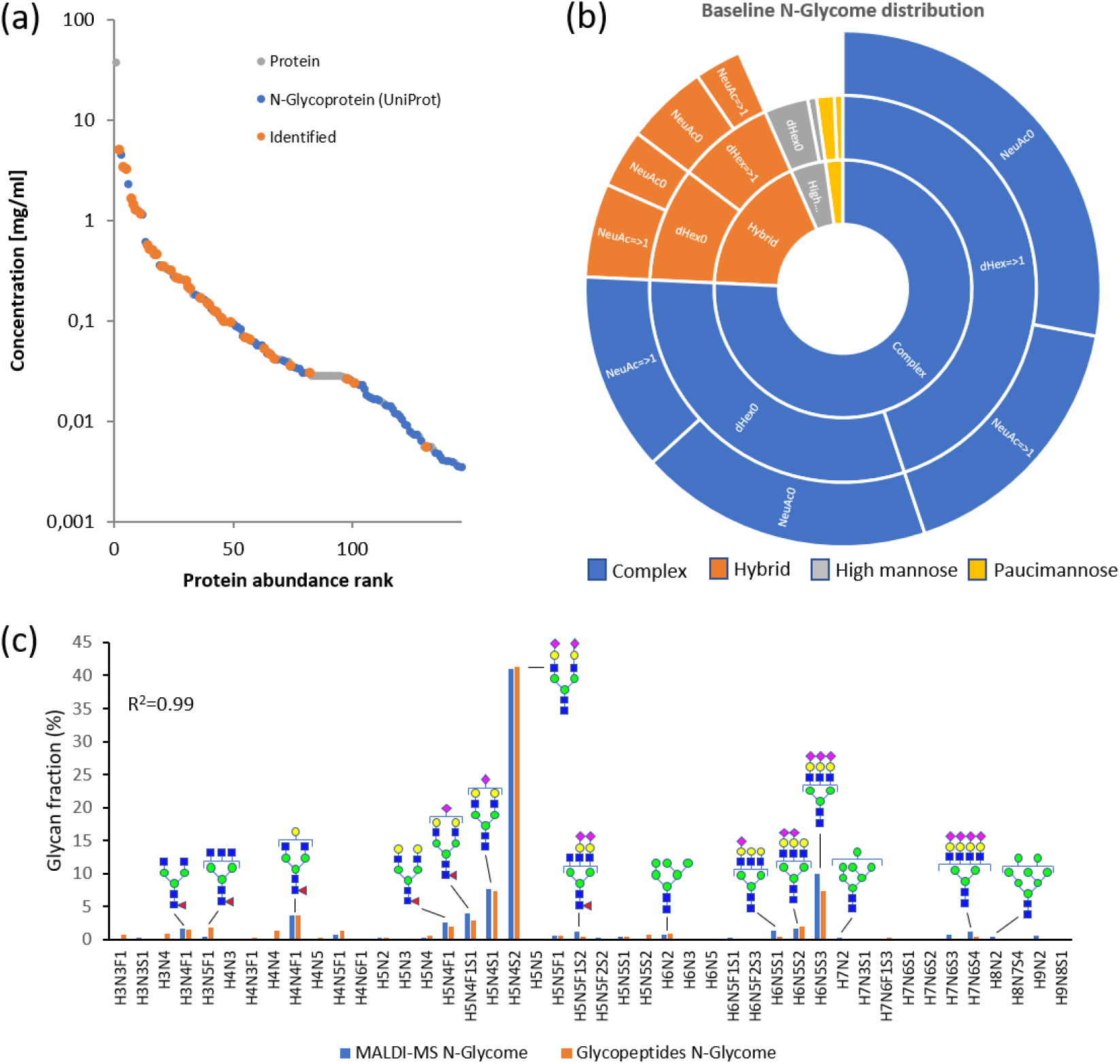
Blood plasma glycoproteome in healthy subjects. **(a)** Glycoprotein abundance distribution of identified glycopeptides. **(b)** Qualitative distribution of identified glycan moieties over major N-glycan classes and traits. **(c)** Relative glycome representations of our experimental glycoproteomics data and glycomics reference data from literature^4^. High abundant glycans are annotated with proposed glycan structures.

Previous clinical glycosylation studies are commonly based on a glycomics workflow in which glycans are released from circulating blood glycoproteins whereas here we analyse intact glycopeptides. To investigate the correlation between the two approaches, we compared our glycoproteomics results for healthy individuals as inferred glycomes with available glycomics data^4^ in figure 2c. The distributions of glycan intensities, expressed as relative glycan fractions, are nearly identical between our glycoproteomics data and reference glycomics dataset (R^2^=0.99) indicating that our workflow has no particular bias towards specific glycoforms and that we can expand current glycomics-based knowledge using our more in-depth glycoproteomics approach.

### Baseline glycoproteome in healthy subjects

The baseline plasma glycoproteome was first assessed by determining the proteome-wide microheterogeneity of glycan-peptide compositions in samples from healthy individuals. The chord diagram in figure 3a depicts the connection of identified glycans to their peptide moiety backbones, indicating a considerable diversity in the number and compositions of glycans between protein glycosylation sites. Both the diversity and partial overlap in glycoforms between individual sites underlines the importance of capturing many complementary site-specific glycosylation profiles within a single measurement for clinical applications to monitor and understand disease-specific glycobiology. Further analysis of glycosylation sites at a higher hierarchical level of glycan traits (figure 3b) showed three distinct clusters where the vast majority of N-glycosylation sites belong to the red cluster that is defined by complex iso-sialylated diantennary glycans. N-glycosylation sites in the blue cluster are decorated with highly fucosylated complex glycans that lack galactose whereas the green cluster is characterized by high mannose glycans. Principle component analysis (PCA) of site-specific glycoform (figure 3c) and glycan trait profiles (figure 3d) revealed consistent glycosylation profiles among healthy individuals with no separation between sample groups based on age- or sex-related differences within the first 10 principal components, respectively. The baseline plasma glycoproteome in healthy individuals thus presents a stable and rich source of reference data to study glycosylation changes in disease.

**Figure 3:**
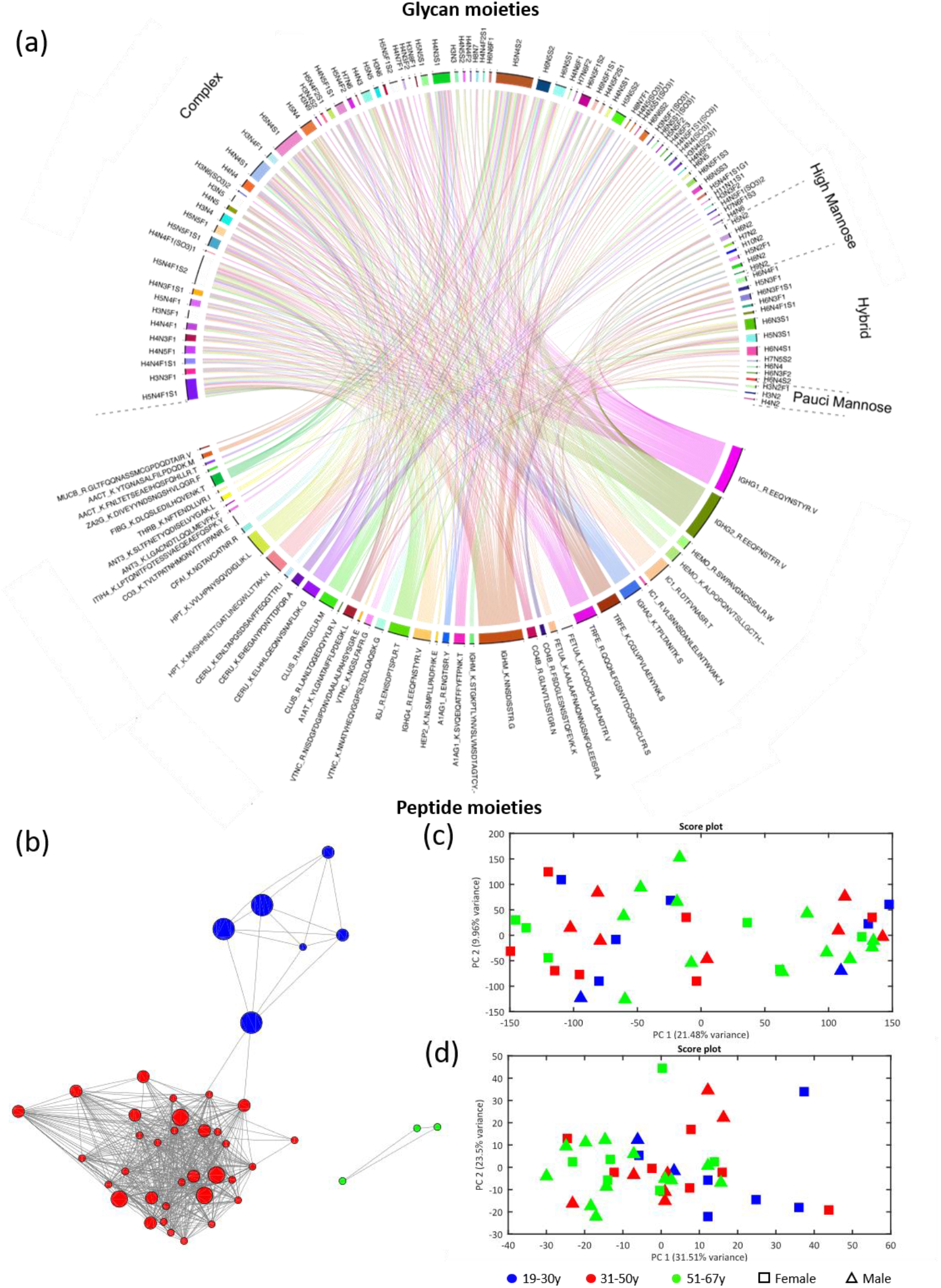
Observed glycoproteome in healthy subjects. **(a)** Chord diagram that visualizes qualitative glycan – peptide relationships of the baseline glycoproteome. Peptides are indexed at the bottom of the diagram and connected via chords to respective identified glycan moieties at the top. **(b)** Glycosylation site similarity network with peptide moiety nodes and edges representing r≥0.8 PCC based on relative glycan trait profiles. Colors are used to indicate distinct clusters. PCA score plots from the first two principal components using **(c)** site-specific glycoform fractions and **(d)** site-specific glycan trait profiles with age classes indicated by color coding and sex by symbols.

### Glycopeptide biomarker discovery in clinical samples

The diversity in glycosylation sites and glycoforms captured by a single glycoproteomics experiment has the potential to identify high-specificity signatures for a variety of human diseases. Indeed, multiple individual glycoproteins in this dataset have already been described as biomarkers for diseases such as cancer, immune disease, cirrhosis and rheumatoid arthritis^33^. Here, we evaluated this added potential of our holistic glycoproteomics workflow using rare human genetic diseases as unique model system. We included genetic defects in a broad range of underlying biological pathways such as glycosyltransferases, sugar metabolism and Golgi homeostasis, of which at least 3 patient samples were available for analysis (10 disease groups). Samples were characterised by intact transferrin glycoprofiling using a clinically validated test by high-resolution mass spectrometry (TRFE IP-MS)^34^. Available data was used to assess the clinical validity of glycopeptide profiling data by comparison with transferrin glycopeptide results at glycome level, showing strong correlation for individual samples (PPC r=0.97; supplementary data).

To identify glycopeptide differentials as possible biomarkers, exploratory chemometrics was performed by PCA using glycopeptide feature data from the 10 CDG groups and controls (n=40). Both glycoproteomics and TRFE IP-MS data showed clear separation between healthy individuals and each of the CDG defects by their first principal component in PCA score plots (figure 4a and supplementary figure 3). Glycopeptide profiling data achieved complete separation between four disease groups (COG5, DYM, NANS and PGM1) whereas the 95% confidence intervals overlapped for TRFE IP-MS data. We also included negative control subsets in which the set of 40 control samples were split into balanced groups of 20 vs 20 samples and unbalanced groups of 35 vs 5 samples. No separation was observed between balanced (20 vs 20) and unbalanced (35 vs 5) control groups as expected. Supervised learning by Partial Least Squares – Discriminant Analysis (PLS-DA)^35^ showed unambiguous disease classification for all patients suffering from any of the 10 CDG defects with Area Under the receiver operator Curve (AUC) values of 1.00 (figures 4a & b and supplementary figure 4). The PLS-DA models achieved better performance indicators for glycoproteomics data over TRFE IP-MS data with respect to AUC for 6 out of 10 defects and Z-score for 8 out of 10 defects. Again, no significant class separation was observed between balanced and unbalanced subsets of control samples. PCA score plots of respectively 82 up to 1542 discriminant features selected by the PLS-DA models showed clear separation between sample classes by their first principal components (PC1) which confirms that the selected features contain strong differentials.

**Figure 4.**
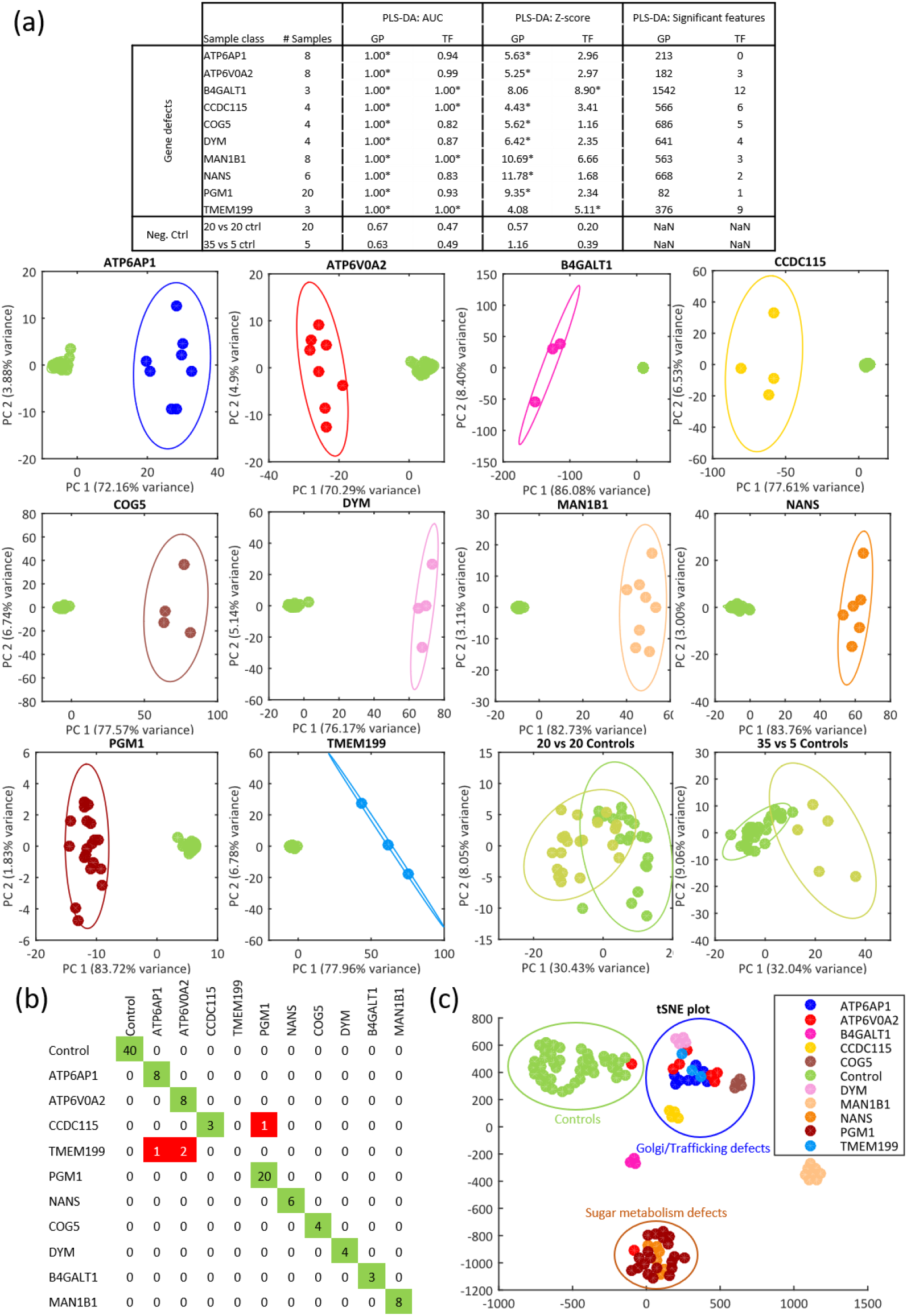
Sample stratification and biomarker identification results for CDG defects. **(a)** Supervised PLS-DA results for glycoproteomics and intact TRFE IP-MS data of CDG defects versus controls as reported area under the curve (AUC), Z-score and number of significant features. **(b)** GA-RF confusion matrix showing the number of correct classifications for samples (rows) to respective genetic defect classes (columns) in green or incorrect classification results in red. **(c)** Unsupervised t-stochastic neighbour embedding plot of GA-RF selected features shows clear separation of samples according to disturbed biological processes. Perturbated Golgi function: ATP6AP1, ATP6V0A2, CCDC115, TMEM199, DYM, COG5. Impaired sugar metabolism: PGM1 and NANS. Independent clusters of N-glycan synthesis enzymes: MAN1B1 and B4GALT1.

Subsequently, we challenged the potential of glycopeptide biomarkers to stratify patients at the level of affected individual genes using a Genetic Algorithm – Random Forest (GA-RF) supervised learning model^36^. Results in figure 4b show that 104 out of 108 samples could be successfully classified to their respective sample groups with high model performance indicators (AUC=0.94 and F1 score=0.88). Three out of four misclassifications were caused by misassignment of TMEM199 samples to similar defects in the V-ATPase complex (ATP6AP1 and ATP6V0A2), that functionally all lead to impairment of the V-ATPase. Unsupervised t-stochastic neighbour embedding (tSNE^37^) of the GA-RF selected features was performed to visualize sample relationships (figure 4c). The tSNE plot shows separated sample clusters of healthy individuals (controls), defects in sugar metabolism (PGM1 and NANS), galactosyltransferase deficiency (B4GALT1), mannosidase deficiency (MAN1B1) and overlapping defects in Golgi homeostasis (ATP6AP1, ATP6V0A2, TMEM199, DYM, CCDC115, and COG5), indicating that the plasma glycoproteomics data can differentiate biologically distinct mechanisms underlying CDG.

### From disease signatures to understanding site-specific glycosylation effects

After the holistic plasma glycoproteomics analyses, we zoomed in to the level of site-specific protein glycosylation to determine if and how these defined genetic defects influence glycosylation in a site-specific manner. Average changes at glycan class level were first determined for five CDG defects in defined steps of the N-glycosylation pathway (figure 5a). Our site-specific data showed that the most prominent glycosylation abnormalities functionally reflect disease mechanisms for each CDG. Reduced fucosylated glycans (F) were observed due to a defect in GDP-fucose transporter SLC35C1, reduced sialylated glycans (Si) and increased hyposialylated glycans (Sh) due to a defect in CMP-sialic acid transporter SLC35A1, while a reduction in GlcNAc-lacking glycans (GN) was observed due to defective UDP-GlcNAc transporter SLC35A3. Additionally, defects in glycan processing enzymes mannosidase MAN1B1 and galactosyltransferase B4GALT1 resulted in increased hybrid structures (H) or galactose-lacking glycans (G), respectively, directly corresponding to the defective step in the N-glycosylation pathway. Subsequent analysis of site-specific glycoforms relative to baseline glycosylation showed that also changes in glycan stoichiometry and expression of disease-specific glycans are directly linked to these respective CDGs (figures 5b & c). We conclude that meaningful site-specific glycosylation changes in disease can reliably be retrieved from holistic plasma glycoproteomics data and correlate with underlying glycobiological mechanisms, demonstrating its potential to translate biomarker signatures to underlying biology.

**Figure 5.**
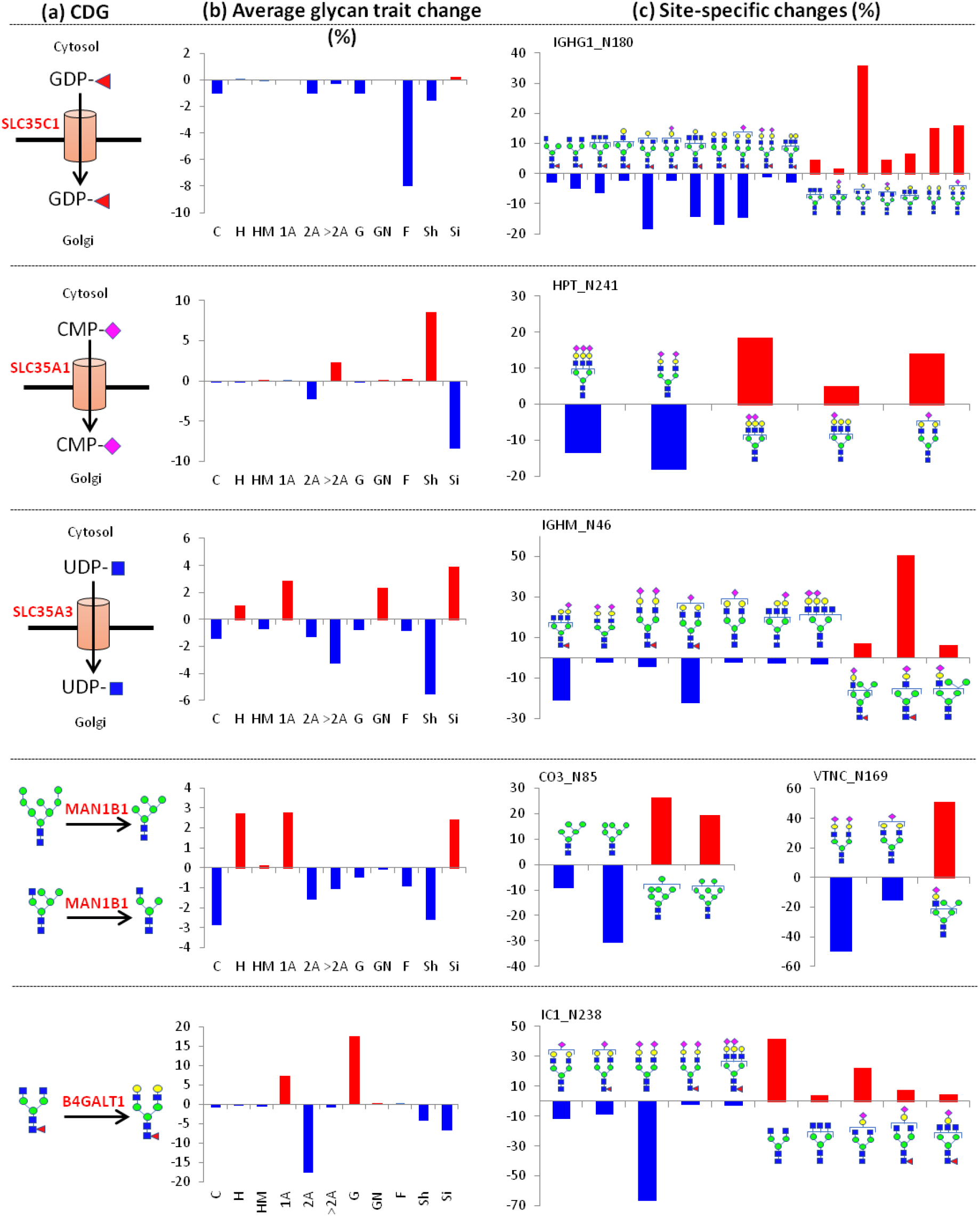
Clinical relevance of site-specific glycosylation changes in CDG. **(a)** Site-specific glycosylation profiles for five CDG types with defined defects in N-glycan biosynthesis were compared to healthy controls, expressed as **(b)** average change in glycan trait distribution. This indicates the average relative change in glycan traits in patients compared to controls from all glycosylation sites. Glycan classes were categorized as follows: C: complex, H: hybrid, HM: high mannose, 1A: single antenna, 2A: two antenna, >2A: 3 or more antenna, G: galactose lacking, GN: GlcNAc lacking, F: fucosylated, Sh: hypo-sialylated, Si: iso-sialylated. **(c)** Illustrative case examples of relative site-specific glycan changes in patient samples from the five CDG defects versus healthy controls, visualized for the indicated sites of glycoproteins immunoglobulin heavy constant gamma 1 (IGHG1), haptoglobin (HPT), immunoglobulin heavy constant mu (IGHM), complement C3 (CO3), vitronectin (VTNC) and plasma protease C1 inhibitor (IC1).

We subsequently aimed to evaluate disease-specific glycosylation changes in an integral analysis that captures system-wide relationships, rather than comparisons of individual glycosylation sites. We therefore visualized glycosylation differences relative to baseline glycosylation in differential chord diagrams (figure 6a, supplementary figure 5). This visualization provided a bird’s eye view of global and site-specific glycoproteome changes from both the glycan and peptide perspectives in a single figure. Clear protein- and site-specific glycosylation differences were observed that correlated strongly with the presence or absence of glycans at respective sites in controls. For example, strongly reduced glycan fucosylation could be readily observed for SLC35C1-CDG on immunoglobulins that possess high levels of fucosylation (about 80-90%) in healthy individuals. In contrast, loss of fucosylation on liver-derived transferrin is barely noticeable due to its very low fucosylation level of ^~^1% in controls. As a second example, the high-mannose glycans of the Asn85 site of complement component CO3 are affected by reduced mannosidase activity in MAN1B1-CDG but remain unaffected in CDG types that impair post-mannose trimming steps in N-glycan biosynthesis (SLC35A1, SLC35A3, SLC35C1 and B4GALT1). As such, baseline glycosylation is a pre-determining factor that dictates if site-specific glycosylation can be affected in specific CDG and thus explains in part protein or site-specific glycosylation changes in disease.

**Figure 6.**
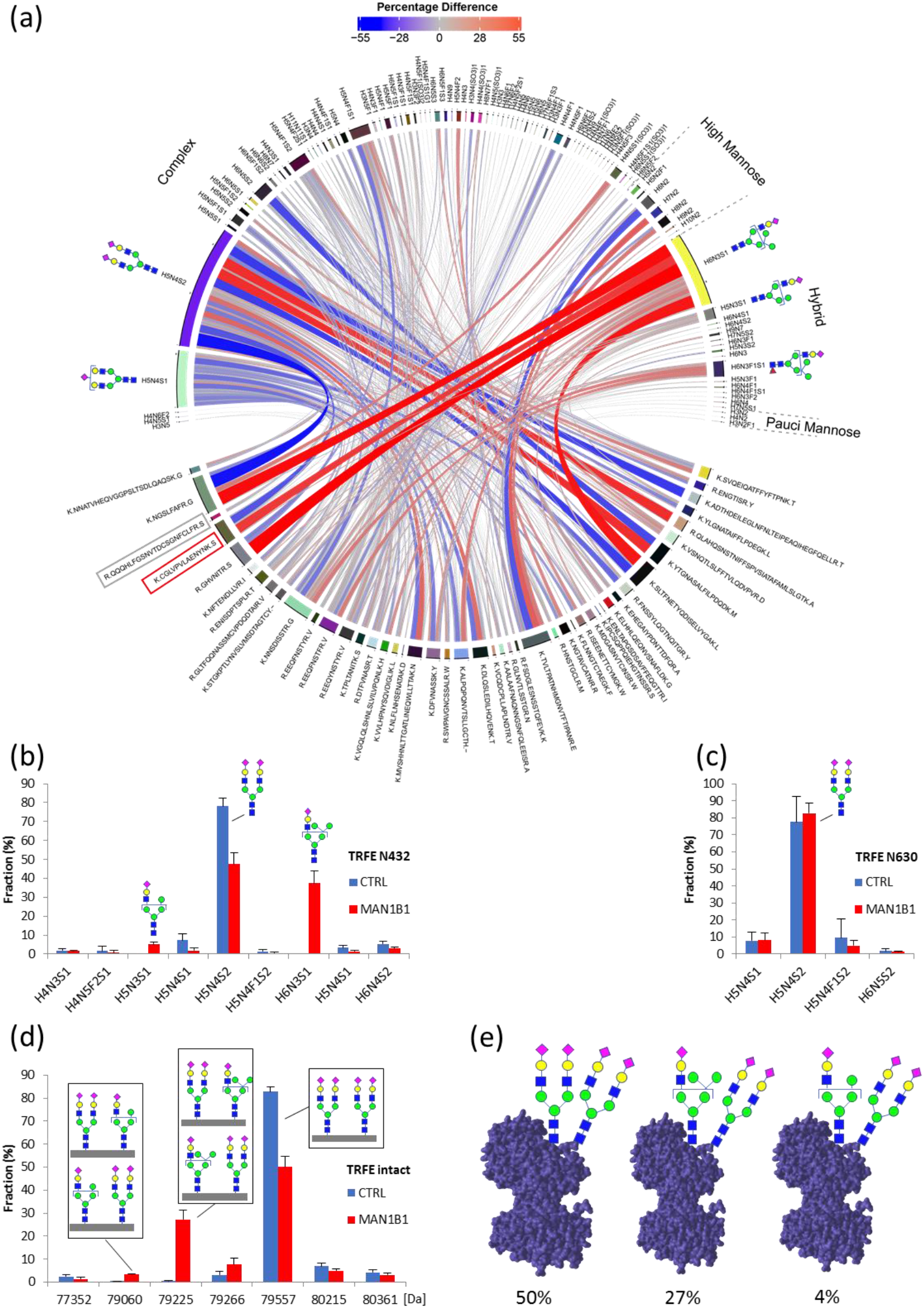
Site-specific glycosylation changes in MAN1B1 deficiency. **(a)** Differential chord diagram depicting all site-specific glycosylation changes relative to baseline glycosylation for MAN1B1 deficiency (see supplementary figure 5 for other CDG types). Chord colors and width indicate relative changes in patients versus controls. TRFE glycopeptides N432 (CGLVPVLAENYNK) and N630 (QQQHLFGSNVTDCSGNFCLFR) are boxed in the chord diagram. Site-specific glycosylation profiles (microheterogeneity) of TRFE at **(b)** Asn432 and **(c)** Asn630 in healthy individuals (n=40) and MAN1B1 deficiency patients (n=8) show that glycosylation of TRFE is exclusively affected at Asn432. **(d)** Macroheterogeneity profiles of TRFE determined in healthy (n=40) and disease subjects (n=8) by intact protein LC-MS show that only one of both glycosylation sites of TRFE is always affected by MAN1B1 deficiency. **(e)** Inferred TRFE glycoform distributions from combining micro- and macro-heterogeneity data visualized with glycan positions indicated in the 3D surface structure of TRFE (pdb: 6JAS).

To investigate site-specific glycosylation changes in more detail we focused on the two glycosylation sites of transferrin (N432 and N630) that have nearly identical baseline glycosylation profiles (complex iso-sialylated diantennary N-glycans) to rule out differences in baseline glycosylation and (tissue-specific) protein synthesis as potential factors. Out of the five genetic defects, only MAN1B1 and SLC35A3 deficiency primarily affected the glycosylation profile at the N432 site, while leaving N630 unaffected (supplementary figure 5, figures 6b and 6c for MAN1B1). For MAN1B1-CDG, this site-specific effect could be confirmed by including macroheterogeneity information (figure 6d). Data from intact TRFE IP MS experiments show that any of the aberrant TRFE glycoforms (H5N4S2-H6N3S1 and H5N4S2-H5N3S1) always contains one normal complex glycan (H5N4S2) and one disease-specific hybrid glycan (H6N3S1 or H5N3S1). Combining both macro- and micro-heterogeneity data enabled a comprehensive view for the exact distribution of TRFE glycoforms in MAN1B1-CDG (figure 6e) where only Asn432 glycosylation is affected. This site-specific effect might be explained by steric hindrance for N-glycosylation enzymes to reach the N432 site, which is located within a pocket of the 3D surface structure of TRFE. This is in line with our observations that N432 is more severely affected than N630 in most CDGs. In general, these results underline the importance of site-specific glycosylation data to explore the glycobiological mechanisms underlying human disease and may provide opportunities to discern genetic defects with shared disease-glycoforms by their site-specific patterns.

## Discussion

In this work we demonstrated the potential of plasma glycoproteomics for patient stratification in clinical studies by virtue of protein- and site-specific glycosylation changes in disease. The analytical robustness of the platform and the strong correlation between baseline glycoproteome and reference glycomes fulfil the prerequisites to diagnose disease through comparative glycoproteomics. Application to a clinical cohort of genetically defined glycosylation diseases (CDG) as proof-of-concept study showed that plasma glycoproteomics is able to detect site-specific glycosylation changes that are directly related to the underlying glycobiology, thereby confirming the clinical relevance of glycoproteome differentials. Our methodology revealed site-specific glycosylation profiles, indicating that the glycosylation status of N-glycoproteins can be affected in a protein- and even site-specific manner in human disease.

The observed heterogeneous protein glycosylation abnormalities in our CDG cohort illustrates the complexity to interpret glycosylation changes in clinical samples. It emphasizes the need to examine protein glycosylation for multiple proteins simultaneously in a site-specific manner using holistic multiplex biomarker analysis methods. We here demonstrated a big step forward in the analysis of a large number of glycopeptides in a single experiment for improved sample stratification as compared to the use of a single glycoprotein biomarker such as transferrin, as used in current CDG diagnostics. Previously observed glycomics changes in various human diseases point towards the existence of novel specific glycoprotein biomarkers in plasma. Here, based on results for CDG patients, we expect that glycopeptide profiling could achieve unparalleled sample stratification for common diseases by providing large-scale site-specific glycosylation data.

At the single protein molecule level, the site-specific behaviour of individual N-glycosylation sites as observed here in CDG patients stresses the importance to take micro-heterogeneity into account next to macro-heterogeneity (unique combinations of glycans at different sites of the same protein molecule^38^), meta-heterogeneity (variation in glycosylation across multiple sites of a given protein^9^) and likely even proteoforms (unique combinations of polypeptide backbone and all present post-translational modifications^39^). Our integrated analysis of transferrin glycopeptides and intact protein mass spectrometry highlights the strength to combine micro- and macroheterogeneity data. It will be key to combine complementary proteomics technologies that characterize proteins from the fragmented bottom-up as well as intact top-down or native perspectives with advanced data modelling as exciting next steps towards improved molecular understanding of glycoproteins and underlying mechanisms in disease.

At the glycoproteome level, the current challenges to translate the potential of glycoproteomics to clinical applications are informatics solutions to reduce, visualize and interpret the complex site-specific glycosylation data with all its intricate relationships. The chord diagrams proposed in this work provide a bird’s eye view of glycoproteome changes in disease but might be of limited practical use when information density increases further. This prompts the development of novel bioinformatic solutions to unravel meaningful correlations between multidimensional glycoproteome changes and clinical phenome data. Successful application of such glycoproteome centric approaches would benefit from recent hardware and software developments for increased glycoproteome coverage and large-scale elucidation of glycan-moiety structures^10^ as additional layer of glycobiology information. In conclusion, glycoproteomics methodologies are emerging to the level of clinical applications in diagnostics and patient stratification, while keeping a detailed level of glycobiological understanding.

## Materials and Methods

### Samples

Plasma samples of patients were obtained from the diagnostic archive of the Radboud University Medical Center, Translational Metabolic Laboratory, expertise center on Congenital Disorders of Glycosylation and used in accordance with Helsinki’s Declaration under local ethical approval (nr 2019-5591). Plasma samples of 40 healthy controls were received from the Sanquin blood bank (Nijmegen, Netherlands) according to their protocols of informed consent. An overview of all samples with associated metadata is available from the supplementary data file.

### Sample preparation for glycopeptide analysis

Ten microliter of plasma was denatured in 10μl urea (8 M urea, 10 mM Tris-HCl pH 8.0) and reduced with 15 μl 10 mM dithiothreitol for 30 min at room temperature (RT). Reduced cysteines were alkylated through incubation with 15 μl 50 mM 2-chloroacetamide (CAA) in the dark for 20 min at RT. Next, proteins were subjected to LysC digestion (1 μg LysC/50μg protein) by incubating the sample at RT for 3 hours. Then, samples were diluted with 3 volumes of 50 mM ammonium bicarbonate and trypsin was added (1 μg trypsin /50 μg protein) for overnight digestion at 37°C. Glycopeptides were enriched using 100 μl Sepharose CL-4B beads slurry (Sigma) per sample well in a 0.20 μm pore size 96 multi well filter plate (AcroPrep Advance, VWR). The beads were washed three times with 20% ethanol and 83% acetonitrile (ACN), respectively, prior to sample application. The sample was then incubated on the beads for 20 min at room temperature on a shaking plate. The filter plate was then centrifuged and beads were first washed three times with 83% ACN and then three times with 83% ACN with 0.1% trifluoroacetic acid (TFA). Next, glycopeptide eluates were collected by incubation of the beads with 50 μl milliQ water for 5 min at room temperature, followed by centrifugation.

### Glycopeptide analysis by LC-MS/MS

Samples were analyzed in randomized order by means of liquid chromatography (nano-Advance, Bruker Daltonics) with online tandem mass spectrometry (maXis Plus, Bruker Daltonics) interfaced via a vacuum-assisted axial desolvation nanoflow electrospray ionization source (CaptiveSpray, Bruker Daltonics) for data-dependent acquisition. Five microliters of sample were loaded onto the trapping column (Acclaim PepMap RSLC, 100 μm × 2 cm, nanoViper, 5 μm 100Å C 18 particles, Thermo Scientific) using 0.1% acetic acid at a flow rate of 7000 nl/min for 3 minutes at RT. Next, peptides were separated on a C18 reversed phase 15 cm length × 7 μm internal diameter C18RP analytical column (Acclaim PepMap RSLC, C18 ReproSil AQ, 1.9 μm particles, 120 Å pore size, Thermo Scientific) at 45°C using a linear gradient of 3–45% ACN 0.1% acetic acid in 60 minutes at a flow rate of 500 nl/min. Electrospray ionization conditions were 3 L/min 180°C N_2_ drying gas, 1500 V capillary voltage and 0.2 Bar N_2_ nebulizer gas flow for gas phase supercharging using ACN as dopant (nanoBooster, Bruker Daltonics). All samples were analysed by data dependent acquisition (AutoMSn) for feature map data generation and glycan-moiety identifications by using a 3s duty cycle at 2 Hz acquisition rate for full MS spectra and a variable number of MS/MS experiments at precursor intensity scaled acquisition rate (4 Hz MS/MS spectral rate at 10.000 counts, 9 Hz MS/MS spectral rate at 75.000 counts). Collision induced dissociation (CID) parameters used for glycan fragmentation: Charge state values [z]: 1; 2; 3; 4; 5; 1; 2; 3; 4; 5; 1; 2; 3; 4; 5; 1; 2; 3; 4; 5, Isolation mass values [m/z]: 300; 300; 300; 300; 300; 500; 500; 500; 500; 500; 1000; 1000; 1000; 1000; 1000; 2000; 2000; 2000; 2000; 2000, Isolation width values [Th]: 4; 4; 4; 4; 4; 5; 5; 5; 5; 5; 8; 8; 8; 8; 8; 20; 20; 20; 20; 20, Collision energy values [eV]: 34; 28; 23; 18; 18; 40; 28; 22; 18; 18; 55; 44; 39; 36; 36; 77; 54; 50; 50; 50, Fallback chargestate [z]: 3. Ion optics tuning: 110 μsec transfer time, 23 μsec pre-pulse storage time, 1200 Volt peak-to-peak (Vpp) collision cell radio frequency (RF).

Respective pooled samples of healthy individuals and selected CDG patient groups (ATP6AP1, ATP6V0A2, B4GALT1, CCDC115, COG5, Cohen syndrome, DYM, MAN1B1, NANS, PGM1, SLC10A7, SLC35A2, TMEM199, VMA21) were analysed in random order to generate peptide-moiety identification data using CID energy stepping to acquire glycan- and peptide-moiety fragmentation data within a merged MS/MS spectrum^27^. To this end, fragmentation spectra were recorded at 100% base collision energy for glycan fragmentation and 200% base collision energy for peptide fragmentation in a 3:7 spectra acquisition stoichiometry. Collision cell tuning was respectively stepped for 50% of each CE MS/MS step time interval from 1000 to 1200 Vpp collision cell RF at 64 to 110 μsec transfer time. Intensity dependent MS/MS spectra acquisition rate was scaled between 2Hz at 10.000 counts and 4Hz at 1.000.000 counts. Precursor ions above 700 *m/z* with charge state z=2+ or higher (preferred charge state range of *z*=2+ to *z*=6+) were selected for MS/MS analysis with active exclusion enabled (excluded after one spectrum, released after 0.5 min, reconsidered precursor if current intensity/previous intensity ≥3, smart exclusion disabled). The mass spectrometry proteomics data have been deposited to the ProteomeXchange Consortium via the PRIDE^40^ partner repository with the dataset identifier PXD034214.

### Intact transferrin mass spectrometry

Glycosylation patterns of intact transferrin that are used as a known biochemical for CDG were analysed by the established CDG diagnostics ESI-MS method^34^. Briefly, rabbit antihuman transferrin antibody (DAKO cat. no. A0061) coupled to *N*-hydroxysuccinimidyl (NHS)-activated Sepharose (GE Healthcare) was used to immunocapture transferrin from plasma samples. Neutralized samples were analysed on a microfluidic 6540 HPLC-chip-QTOF instrument (Agilent Technologies) using a reversed phase C8 stationary phase. Data analysis was performed using Agilent Mass Hunter Qualitative Analysis Software B.04.00 in combination with the Agilent BioConfirm Software for charge deconvolution using the maximum entropy algorithm.

### Database searches and consensus feature map generation

Raw data were processed using DataAnalysis 4.2 for post-acquisition internal mass calibration (sodium-acetate clusters) and subsequent export of MS/MS data to MGF file format. Mass calibrated raw data files were converted to mzML using compassXport (Bruker Daltonics) and subsequently processed by OpenMS (Knime v3.2.1) to generate a consensus feature map that contains intensities and associated metadata for aligned features from all analysis files. The OpenMS workflow can be downloaded from proteomeXchange (PXD034214). The MGF-files were processed in ProteinScape™ 3.1 (Bruker Daltonics) to classify glycopeptide fragmentation spectra using sugar mass distance pattern matching and to calculate peptide- and glycan-moiety masses, respectively. Classified glycopeptide fragmentation MS/MS spectra were searched against the Carbbank glycan structure database (date: 2016-08-05) using GlycoQuest (ProteinScape™ 3.1, Bruker Daltonics) and following acceptance criteria: score>20, >10% spectral intensity coverage and >10% theoretical fragment ion coverage (B-,Y- and internal-fragment ions). An *in-house* developed Perl script was used to remove oxonium ions (B-, and i-ions) based on 25% detection frequency and glycan-peptide fragment ions (Y-ions) with mass above the calculated non-glycosylated peptide moiety from CE stepping fragmentation spectra and searched against the human SwissProt protein sequence database (date: 2016-08-05) by MASCOT (v2.5.1.1, Matrix science) with the following settings: tryptic cleavage specificity, precursor mass tolerance of 20.0 ppm, MS/MS tolerance of 0.05 Da, allowing for 1 missed cleavage and a fixed carbamidomethyl modification (C), and variable deamidation (NQ), oxidation (M), acetyl (protein N-terminus), HexNAc (N) and pyro-carbamidomethyl (peptide N-terminus) modifications, with percolator enabled to achieve ≤1.0% false discovery rate.

### Data pre-processing and integration

*In-house* developed MATLAB R2014b functions and Perl scripts were used to map the identification data onto features of the consensus feature map using a rectangular bucketing approach and to perform subsequent data pre-processing and downstream analyses. All functions and scripts used in this work are available from proteomeXchange (PXD034214). Briefly, in pre-processing relevant features within specified retention time and mass-to-charge ranges (10 - 67 min, 600 - 3700 *m/z)* and with detection frequency of at least 75% in samples within any sample group were subjected to quantile intensity normalization prior to missing value imputation for subsequent machine learning approaches. Identified non-redundant peptide- and glycan-moieties from individual search results were first subjected to retention time alignment by LOESS regression and subsequently linked to consensus features in rectangular buckets (1 min retention time width x 20 ppm mass width). Inter-search result conflicts were resolved by majority vote calling for respectively peptide- and glycan-moieties prior to filtering out cases where the sum of the glycan- and peptide-moiety masses was greater than the (combined) feature mass after proximal HexNAc classification ambiguity correction, when possible. Next, identification results from annotated features were transferred onto unannotated co-eluting features with identical charge deconvoluted mass. Finally, peptide identifications from fully annotated features (both peptide- and glycan-moiety elucidated) were matched to co-eluting features with identified glycan-moieties when the calculated peptide-moiety mass from the glycan identification corresponded with the experimental mass of the identified peptide-moiety mass within 20 ppm and number of sialic acids b etween both features’ glycan-moieties were identical.

### Bioinformatics analysis

Site-specific glycosylation profiles were generated as glycoform fractions by dividing the glycan intensity by the total sum of glycan intensities for each respective peptide sequence. Composition Based glycan Classification (CBC) was performed using computational logic described as pseudo algorithm and available from the proteomeXchange deposition. Peptide sequences lacking the consensus [N-X-S/T] N-glycosylation sequon and peptide-moieties with 1 or more missed cleavages were removed from the dataset. Baseline and transferrin glycopeptide-based glycome inference was performed by summing intensities from identical glycoform features and expressed as relative glycan fractions. Glycome inference from intact transferrin – MS macro-heterogeneity data was performed by summing the respective glycan intensity for each protein glycoform after correction for the number of occupied N-glyosylation site positions and expressed as relative glycan fractions. Chord diagrams for glycoproteome visualizations were prepared using the circlize package^41^ in R programming language v4.1.1^42^. To generate a site-specific CBC similarity network we first constructed a Pearson’s correlation matrix from the CBC distributions of all peptide-moieties after which correlations between glycosylation sites with r ≥ 0.8 were used as edges for peptide-moiety nodes in the MATLAB biograph function for network generation.

### Supervised machine learning and nonlinear dimensional reduction

Partial Least Squares - Discriminant Analysis (PLS-DA) models were constructed using a repeated double cross validation procedure in 21 model iterations with number of latent variables optimized in a Leave-One-Out inner cross validation loop and classification models build using a 3-fold outer cross validation loop resulting in predicted class labels. PLS-DA features with variable importance in project (VIP) >1 and with false discovery rate (FDR) <1% based on 2000 permutations were selected as discriminant features from the models. In Genetic Algorithm – Random Forest (GA-RF)^36^ classification a genetic algorithm was used to select significant features using the number of misclassifications from the random forest algorithm as fitness function. GA parameters used were population size: 200, selection function: tournament, uniform mutation: 0.1, crossover heuristic: 0.2, elite count: 2, stall generation limit: 10. Each RF classification was performed using 100 trees with a leave-one-out cross validation for prediction and subsequent performance calculation. Principle Component Analysis (PCA) and mahalanobis distance measures were performed using standard MATLAB R2014b functions. Nonlinear dimensional reduction using t-distributed Stochastic Neighbor Embedding (t-SNE) was performed using the non-parametric t-SNE MATLAB function from van der Maaten *et al* ^37^with 6 initial dimensions and 10 perplexity.

## Acknowledgements

This research was part of the Netherlands X-omics Initiative and partially funded by NWO (project 184.034.019), supported by ZonMw Medium Investment Grant (40-00506-98-9001) and EUROGLYCAN-omics (ERARE18-117), under the frame of E-Rare-3, the ERA-Net for Research on Rare Diseases.

## References

1 Schjoldager, K. T., Narimatsu, Y., Joshi, H. J. & Clausen, H. Global view of human protein glycosylation pathways and functions. Nature reviews. Molecular cell biology 21, 729–749, doi:10.1038/s41580-020-00294-x (2020).

2 Reily, C., Stewart, T. J., Renfrow, M. B. & Novak, J. Glycosylation in health and disease. Nature reviews. Nephrology 15, 346–366, doi:10.1038/s41581-019-0129-4 (2019).

3 Lefeber, D. J. Protein-Specific Glycoprofiling for Patient Diagnostics. Clinical chemistry 62, 9–11, doi:10.1373/clinchem.2015.248518 (2016).

4 Hipgrave Ederveen, A. L., de Haan, N., Baerenfaenger, M., Lefeber, D. J. & Wuhrer, M. Dissecting Total Plasma and Protein-Specific Glycosylation Profiles in Congenital Disorders of Glycosylation. International journal of molecular sciences 21, doi:10.3390/ijms21207635 (2020).

5 Merleev, A. A. et al. A site-specific map of the human plasma glycome and its age and gender-associated alterations. Scientific reports 10, 17505, doi:10.1038/s41598-020-73588-x (2020).

6 Gilgunn, S., Conroy, P. J., Saldova, R., Rudd, P. M. & O’Kennedy, R. J. Aberrant PSA glycosylation--a sweet predictor of prostate cancer. Nature reviews. Urology 10, 99–107, doi:10.1038/nrurol.2012.258 (2013).

7 Narimatsu, H. et al. Current Technologies for Complex Glycoproteomics and Their Applications to Biology/Disease-Driven Glycoproteomics. Journal of proteome research 17, 4097–4112, doi:10.1021/acs.jproteome.8b00515 (2018).

8 Chen, Z., Huang, J. & Li, L. Recent advances in mass spectrometry (MS)-based glycoproteomics in complex biological samples. Trends in analytical chemistry: TRAC 118, 880–892, doi:10.1016/j.trac.2018.10.009 (2019).

9 Caval, T., Heck, A. J. R. & Reiding, K. R. Meta-heterogeneity: evaluating and describing the diversity in glycosylation between sites on the same glycoprotein. Molecular & cellular proteomics: MCP, doi:10.1074/mcp.R120.002093 (2020).

10 Kawahara, R. et al. Community evaluation of glycoproteomics informatics solutions reveals high-performance search strategies for serum glycopeptide analysis. Nature methods 18, 1304–1316, doi:10.1038/s41592-021-01309-x (2021).

11 Hoffmann, M., Marx, K., Reichl, U., Wuhrer, M. & Rapp, E. Site-specific O-Glycosylation Analysis of Human Blood Plasma Proteins. Molecular & cellular proteomics: MCP 15, 624–641, doi:10.1074/mcp.M115.053546 (2016).

12 Yu, Q. et al. Electron-Transfer/Higher-Energy Collision Dissociation (EThcD)-Enabled Intact Glycopeptide/Glycoproteome Characterization. Journal of the American Society for Mass Spectrometry 28, 1751–1764, doi:10.1007/s13361-017-1701-4 (2017).

13 Totten, S. M., Feasley, C. L., Bermudez, A. & Pitteri, S. J. Parallel Comparison of N-Linked Glycopeptide Enrichment Techniques Reveals Extensive Glycoproteomic Analysis of Plasma Enabled by SAX-ERLIC. Journal of proteome research 16, 1249–1260, doi:10.1021/acs.jproteome.6b00849 (2017).

14 Wang, S. et al. Profiling of Endogenously Intact N-Linked and O-Linked Glycopeptides from Human Serum Using an Integrated Platform. Journal of proteome research 19, 1423–1434, doi:10.1021/acs.jproteome.9b00592 (2020).

15 Peng, J. et al. High Anti-Interfering Profiling of Endogenous Glycopeptides for Human Plasma by the Dual-Hydrophilic Metal-Organic Framework. Analytical chemistry 91, 4852–4859, doi:10.1021/acs.analchem.9b00542 (2019).

16 King, S. L. et al. Characterizing the O-glycosylation landscape of human plasma, platelets, and endothelial cells. Blood advances 1, 429–442, doi:10.1182/bloodadvances.2016002121 (2017).

17 Lin, C. H., Krisp, C., Packer, N. H. & Molloy, M. P. Development of a data independent acquisition mass spectrometry workflow to enable glycopeptide analysis without predefined glycan compositional knowledge. Journal of proteomics 172, 68–75, doi:10.1016/j.jprot.2017.10.011 (2018).

18 Saraswat, M. et al. Tongue Cancer Patients Can be Distinguished from Healthy Controls by Specific N-Glycopeptides Found in Serum. Proteomics. Clinical applications 12, e1800061, doi:10.1002/prca.201800061 (2018).

19 Chang, T. T. et al. Plasma proteome plus site-specific N-glycoprofiling for hepatobiliary carcinomas. The journal of pathology. Clinical research 5, 199–212, doi:10.1002/cjp2.136 (2019).

20 Zhang, Y. et al. Comparative Glycoproteomic Profiling of Human Body Fluid between Healthy Controls and Patients with Papillary Thyroid Carcinoma. Journal of proteome research 19, 2539–2552, doi:10.1021/acs.jproteome.9b00672 (2020).

21 Qin, H. et al. Highly Efficient Analysis of Glycoprotein Sialylation in Human Serum by Simultaneous Quantification of Glycosites and Site-Specific Glycoforms. Journal of proteome research 18, 3439–3446, doi:10.1021/acs.jproteome.9b00332 (2019).

22 Zhang, Y. et al. Glyco-CPLL: An Integrated Method for In-Depth and Comprehensive N-Glycoproteome Profiling of Human Plasma. Journal of proteome research 19, 655–666, doi:10.1021/acs.jproteome.9b00557 (2020).

23 Joenvaara, S. et al. Quantitative N-glycoproteomics reveals altered glycosylation levels of various plasma proteins in bloodstream infected patients. PloS one 13, e0195006, doi:10.1371/journal.pone.0195006 (2018).

24 Zhang, Y. et al. Systems analysis of singly and multiply O-glycosylated peptides in the human serum glycoproteome via EThcD and HCD mass spectrometry. Journal of proteomics 170, 14–27, doi:10.1016/j.jprot.2017.09.014 (2018).

25 Deleon-Pennell, K. Y. et al. Glycoproteomic Profiling Provides Candidate Myocardial Infarction Predictors of Later Progression to Heart Failure. ACS omega 4, 1272–1280, doi:10.1021/acsomega.8b02207 (2019).

26 Wada, Y., Tajiri, M. & Yoshida, S. Hydrophilic affinity isolation and MALDI multiple-stage tandem mass spectrometry of glycopeptides for glycoproteomics. Analytical chemistry 76, 6560–6565, doi:10.1021/ac049062o (2004).

27 Hinneburg, H. et al. The Art of Destruction: Optimizing Collision Energies in Quadrupole-Time of Flight (Q-TOF) Instruments for Glycopeptide-Based Glycoproteomics. Journal of the American Society for Mass Spectrometry 27, 507–519, doi:10.1007/s13361-015-1308-6 (2016).

28 Pfeuffer, J. et al. OpenMS - A platform for reproducible analysis of mass spectrometry data. Journal of biotechnology 261, 142–148, doi:10.1016/j.jbiotec.2017.05.016 (2017).

29 Thiele, H., Glandorf, J. & Hufnagel, P. Bioinformatics strategies in life sciences: from data processing and data warehousing to biological knowledge extraction. Journal of integrative bioinformatics 7, 141, doi:10.2390/biecoll-jib-2010-141 (2010).

30 Perkins, D. N., Pappin, D. J., Creasy, D. M. & Cottrell, J. S. Probability-based protein identification by searching sequence databases using mass spectrometry data. Electrophoresis 20, 3551–3567, doi:10.1002/(sici)1522-2683(19991201)20:18<3551::aid-elps3551>3.0.co;2-2 (1999).

31 Alagesan, K. & Kolarich, D. To enrich or not to enrich: Enhancing (glyco)peptide ionization using the Captive Spray nano Booster™. bioRxiv, 597922, doi:10.1101/597922 (2019).

32 Lange, E., Tautenhahn, R., Neumann, S. & Gröpl, C. Critical assessment of alignment procedures for LC-MS proteomics and metabolomics measurements. BMC Bioinformatics 9, 375–375, doi:10.1186/1471-2105-9-375 (2008).

33 Clerc, F. et al. Human plasma protein N-glycosylation. Glycoconjugate journal 33, 309–343, doi:10.1007/s10719-015-9626-2 (2016).

34 van Scherpenzeel, M., Steenbergen, G., Morava, E., Wevers, R. A. & Lefeber, D. J. High-resolution mass spectrometry glycoprofiling of intact transferrin for diagnosis and subtype identification in the congenital disorders of glycosylation. Translational research: the journal of laboratory and clinical medicine 166, 639–649 e631, doi:10.1016/j.trsl.2015.07.005 (2015).

35 Lee, L. C., Liong, C. Y. & Jemain, A. A. Partial least squares-discriminant analysis (PLS-DA) for classification of high-dimensional (HD) data: a review of contemporary practice strategies and knowledge gaps. The Analyst 143, 3526–3539, doi:10.1039/c8an00599k (2018).

36 Paul, D. et al. Feature selection for outcome prediction in oesophageal cancer using genetic algorithm and random forest classifier. Computerized medical imaging and graphics: the official journal of the Computerized Medical Imaging Society 60, 42–49, doi:10.1016/j.compmedimag.2016.12.002 (2017).

37 Laurens van der Maaten, G. H. Visualizing Data using t-SNE. Journal of Machine Learning Research 9, 2579–2605 (2008).

38 Zacchi, L. F. & Schulz, B. L. N-glycoprotein macroheterogeneity: biological implications and proteomic characterization. Glycoconjugate journal 33, 359–376, doi:10.1007/s10719-015-9641-3 (2016).

39 Smith, L. M., Kelleher, N. L. & Consortium for Top Down, P. Proteoform: a single term describing protein complexity. Nature methods 10, 186–187, doi:10.1038/nmeth.2369 (2013).

40 Perez-Riverol, Y. et al. The PRIDE database resources in 2022: a hub for mass spectrometry-based proteomics evidences. Nucleic Acids Res 50, D543–d552, doi:10.1093/nar/gkab1038 (2022).

41 Gu, Z., Gu, L., Eils, R., Schlesner, M. & Brors, B. circlize implements and enhances circular visualization in R. Bioinformatics 30, 2811–2812, doi:10.1093/bioinformatics/btu393 (2014).

42 Team, R. D. C. (Vienna, Austria, 2009).

