## Supplementary Figures for "Plasma glycoproteomics delivers high-specificity disease biomarkers by detecting site-specific glycosylation abnormalities"

### Supplementary figure 1

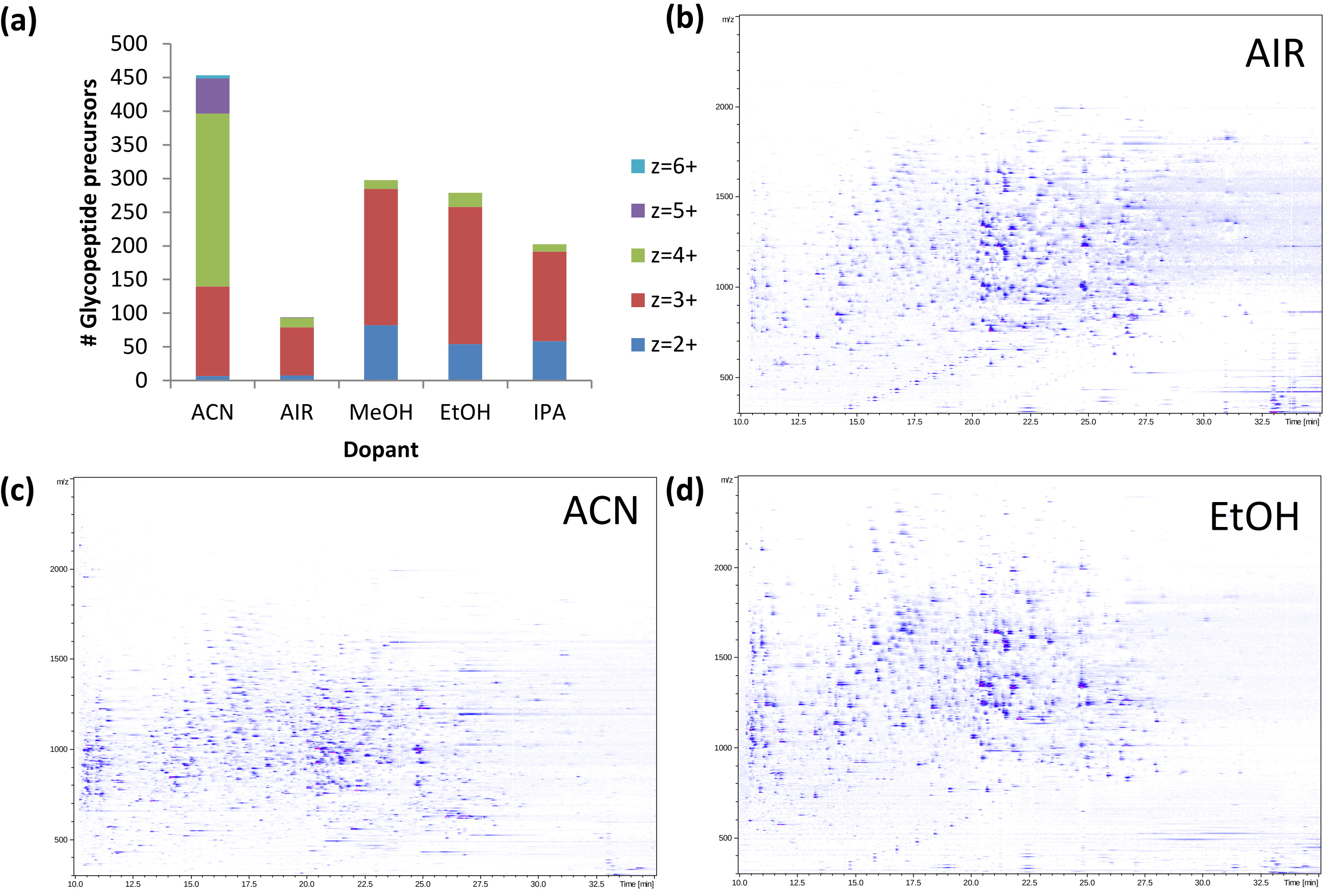


**Supplementary figure 1.** NanoBooster dopant effects on intact glycopeptides enriched from full blood plasma by ZIC-HILIC (ProteoExtract Glycopeptide Enrichment kit pn72103; Merck Millipore). (a) Total number of glycopeptide precursors and according charge state distribution using different nanoBooster dopants. Illustrative ion map data representations of analyses performed using **(b)** filtered Air [default captivesprayer operation without nanoBooster] and using **(c)** acetonitrile or **(d)** ethanol as nanoBooster dopant. Abbreviations: ACN – acetonitrile dopant, AIR – filtered ambient air, ACN – acetonitrile dopant, EtOH – ethanol dopant.

### Supplementary figure 2

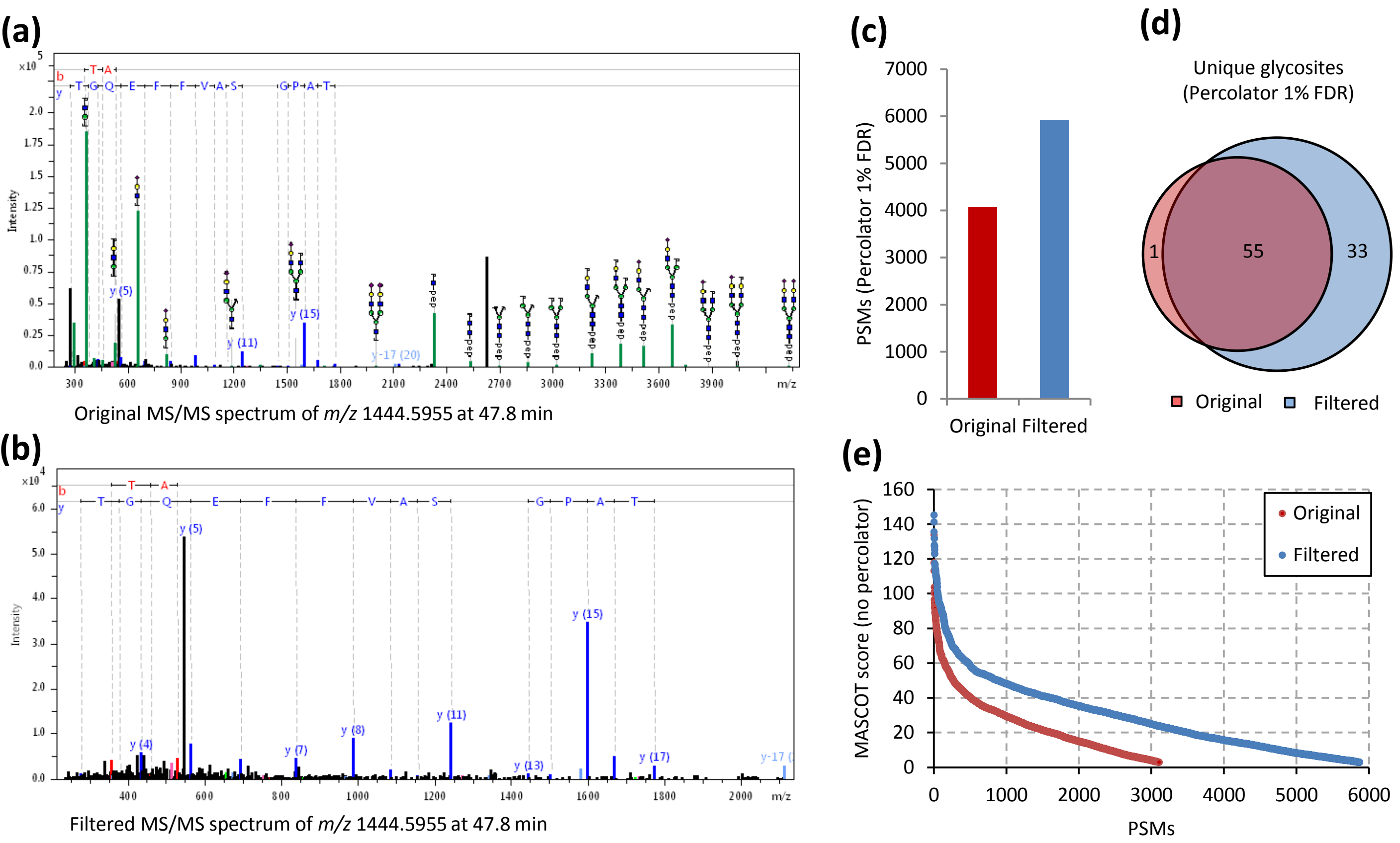


**Supplementary figure 2.** Fragment ion filtering from MS/MS spectra improve MASCOT peptide-moiety database search identifications. **(a)** Original charge deconvoluted MS/MS spectrum with annotated glycan- and peptide-moiety fragments **(b)** Filtered charge deconvoluted MS/MS spectrum with annotated peptide fragment ions **(c)** MS/MS spectrum filtering increases the number of peptide moiety-spectrum matches. **(d)** Venn diagram for unique peptide-moieties identification from original and filtered MS/MS spectra shows significant increase in unique identifications after filtering. The single peptide moiety match unique for original MS/MS spectra was a false positive result as the identified peptide sequence did not contain an N-glycosylation site motif and partly based on glycan B-ion fragments. **(e)** MASCOT ion score index plot of original and filtered MS/MS peptide moiety-spectrum matches (no percolator).

### Supplementary figure 3

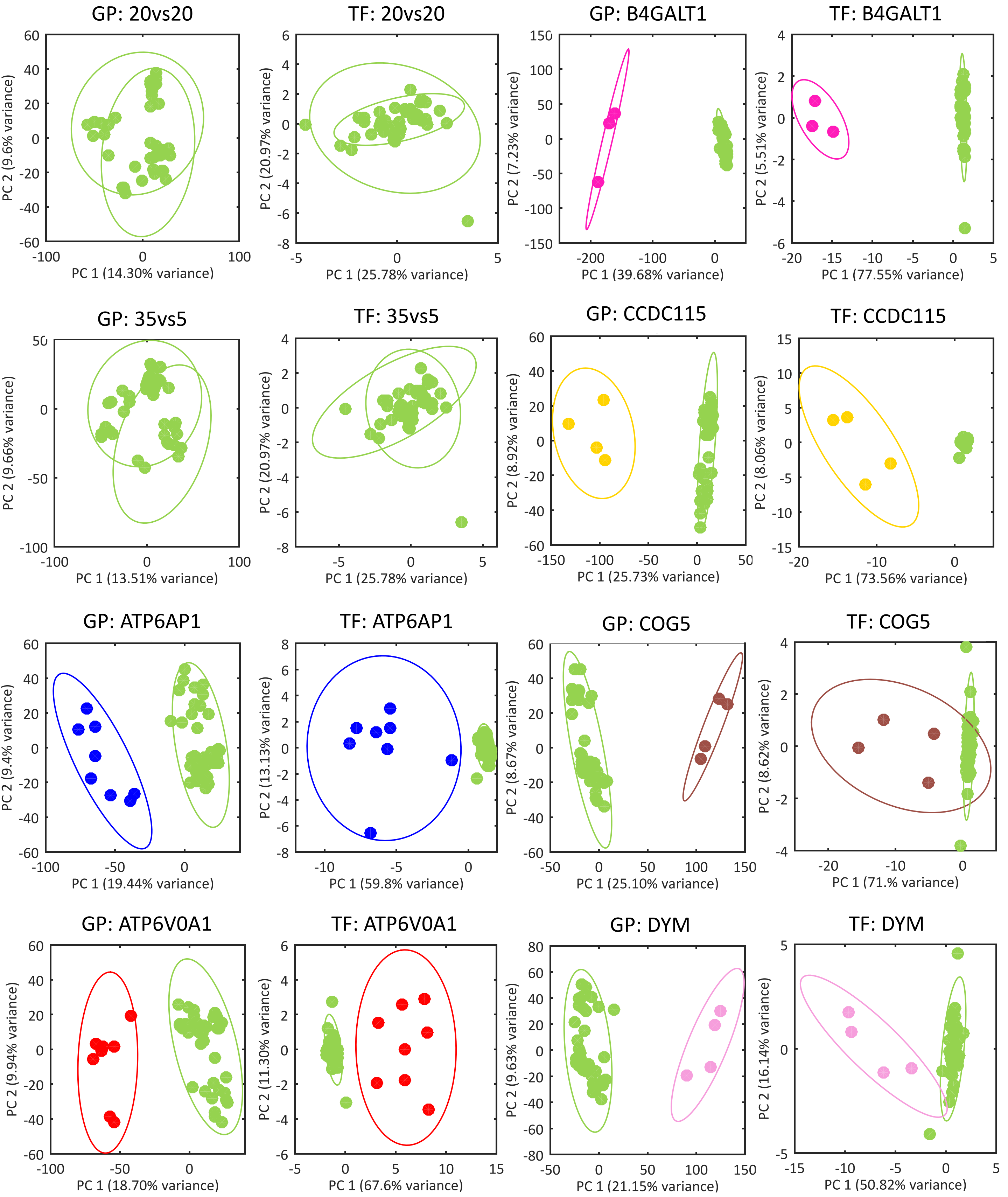

### Supplementary figure 3 - continued

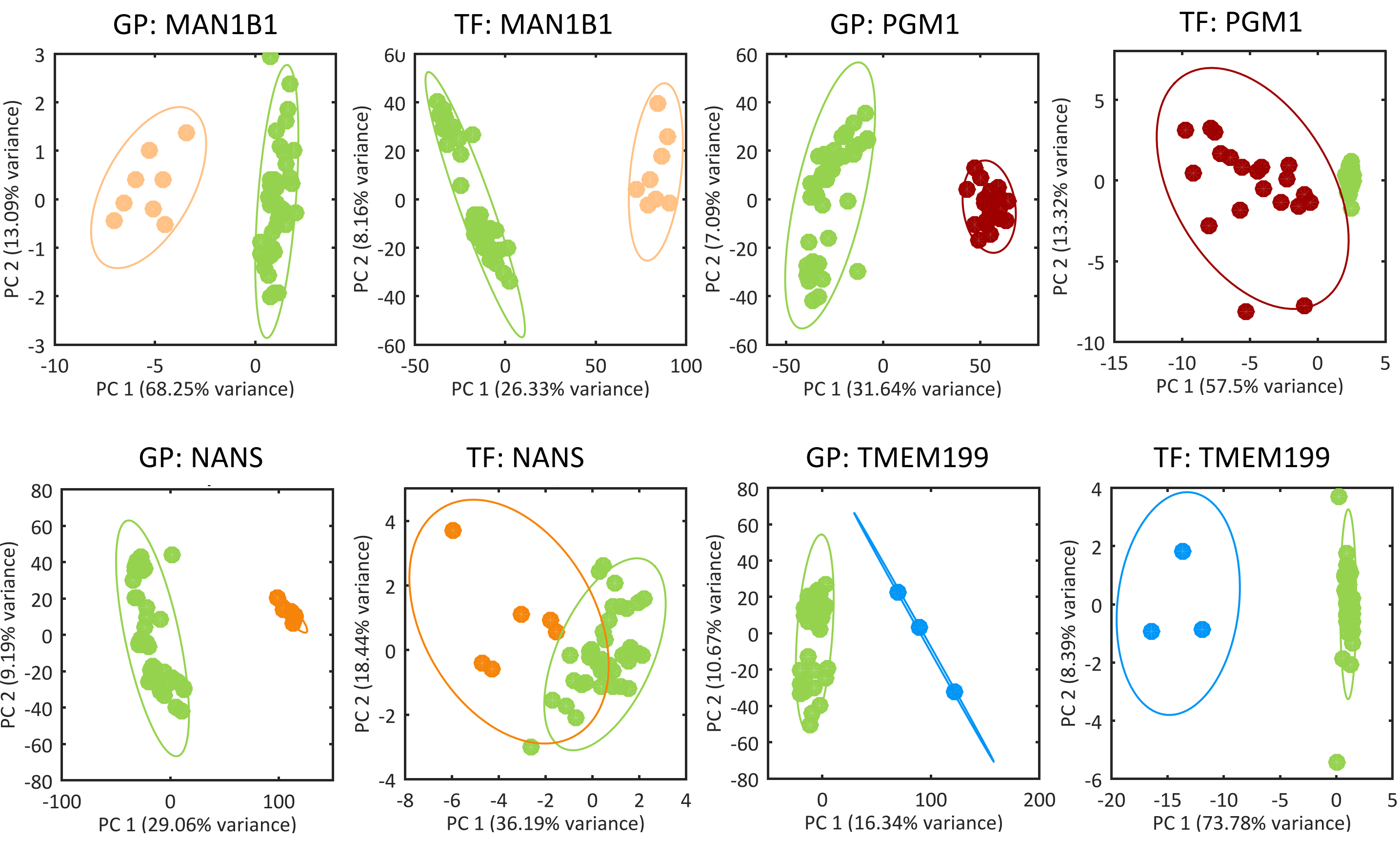


**Supplementary figure 3.** Principle component score plots for glycopeptide profiling- (GP) and intact transferrin LC-MS (TF) data using all available features in binary comparisons of CDGs versus controls. Green dots in these score plots are n=40 healthy individuals. Other colored dots correspond respectively with ATP6AP1 (n=8), ATP6V0A2 (n=8), B4GALT1 (n=3), CCDC115 (n=4), COG5 (n=4), DYM (n=4), MAN1B1 (n=8), NANS (n=6), PGM1 (n=20) and TMEM199 (n=3) CDG patients.

### Supplementary figure 4

**Supplementary figure 4.** PLS-DA models for all binary comparisons of selected CDGs versus controls (n=40) using intact Transferrin LC-MS (TF) and glycopeptide profiling (GP) data. Colored dots show classification result for each of the 21 individual PLS-DA models that were used to generate the final PLS-DA model for each sample, respectively. The black dot shows the classification result for each sample by the final (combined) PLS-DA model. Included CDGs are: ATP6AP1 (n=8), ATP6V0A2 (n=8), B4GALT1 (n=3), CCDC115 (n=4), COG5 (n=4), DYM (n=4), MAN1B1 (n=8), NANS (n=6), PGM1 (n=20) and TMEM199 (n=3).

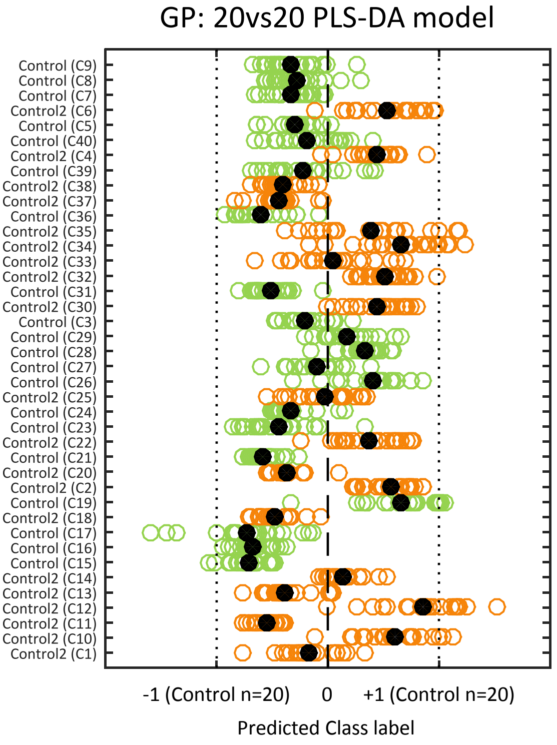


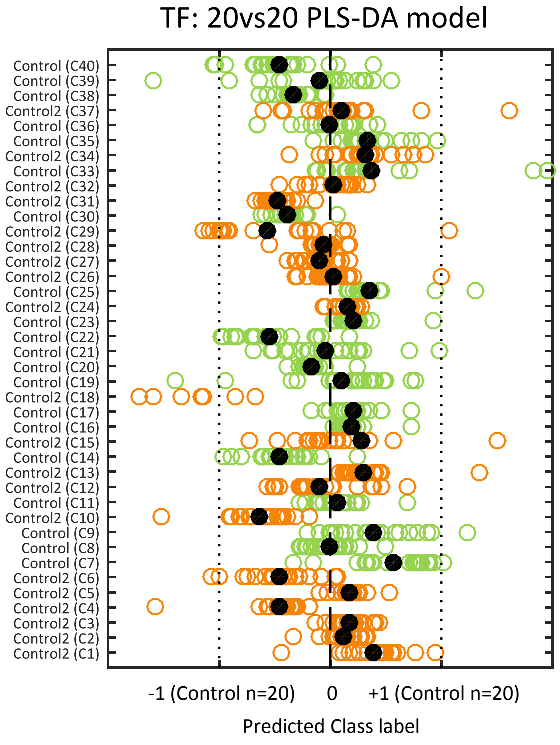


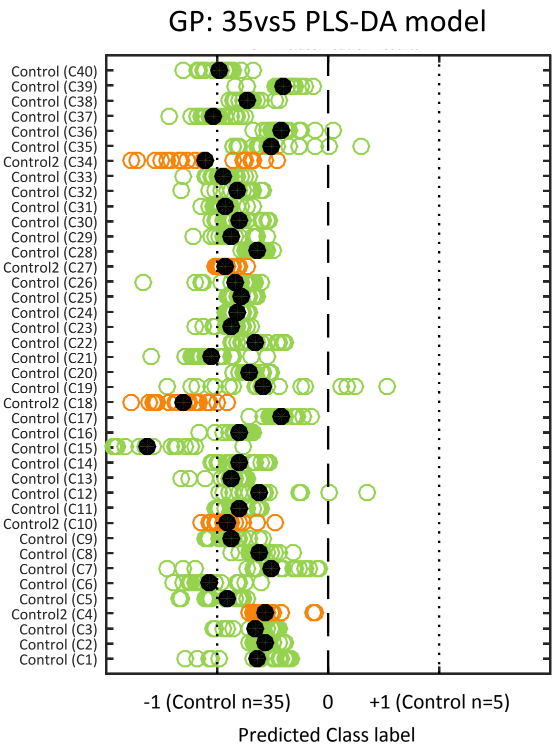


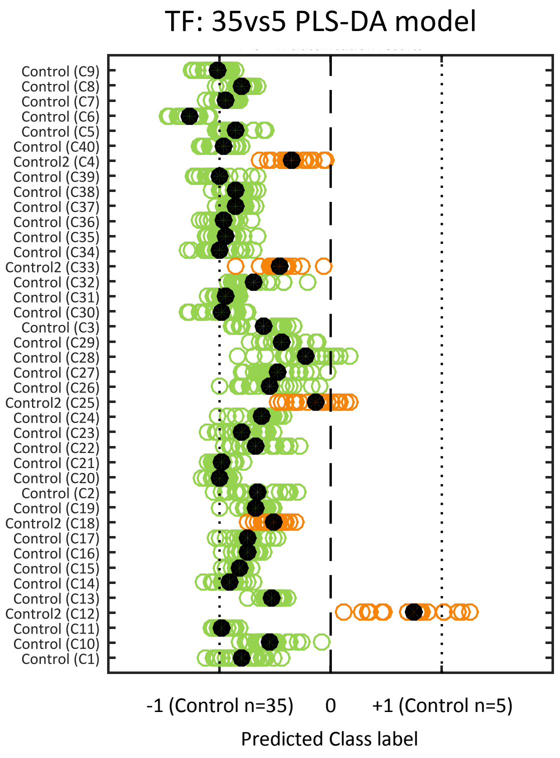

### Supplementary figure 4 - continued

GP: ATP6AP1 PLS-DA model

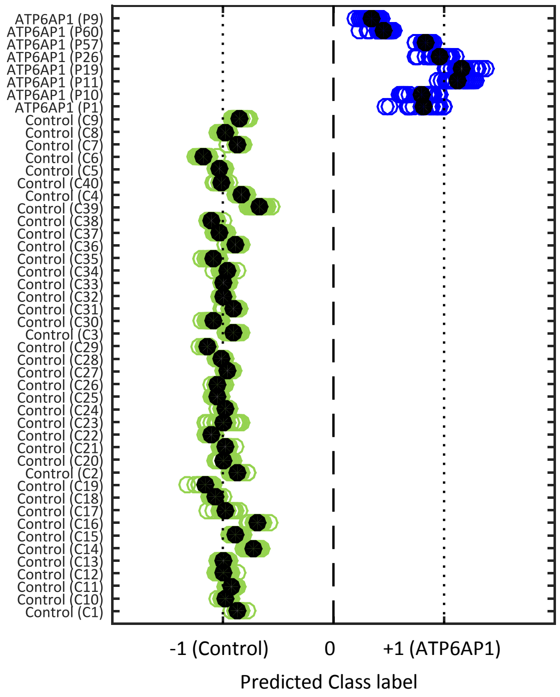


TF: ATP6AP1 PLS-DA model

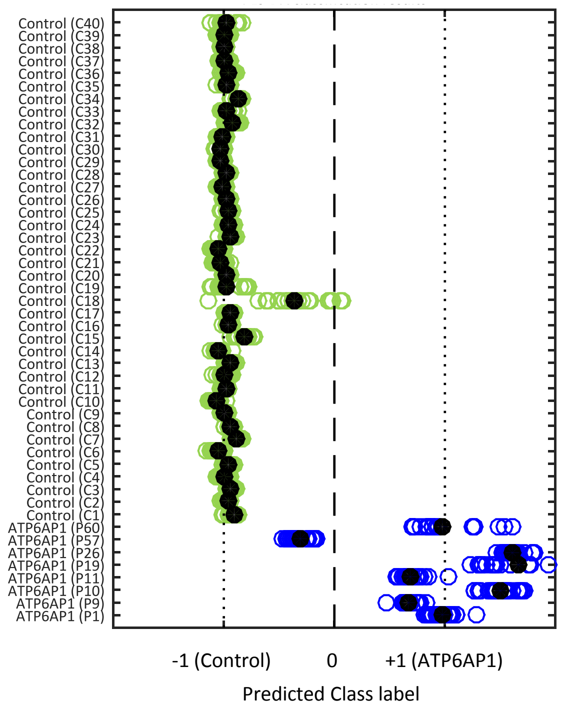


GP: ATP6V0A1 PLS-DA model

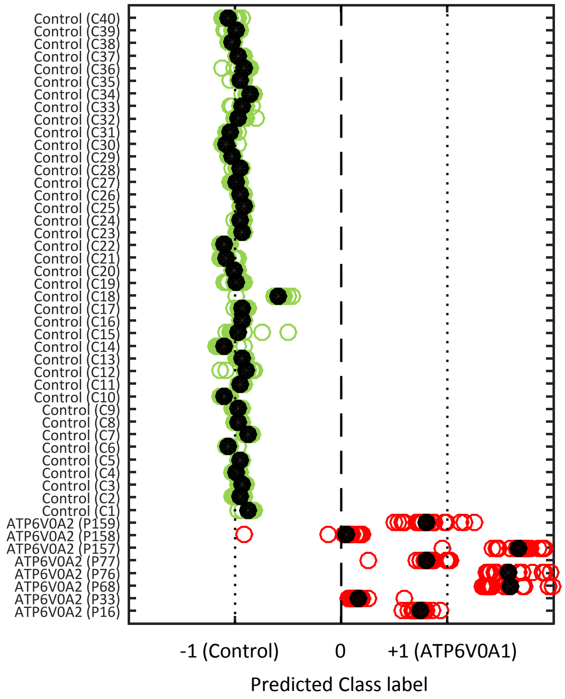


TF: ATP6V0A1 PLS-DA model

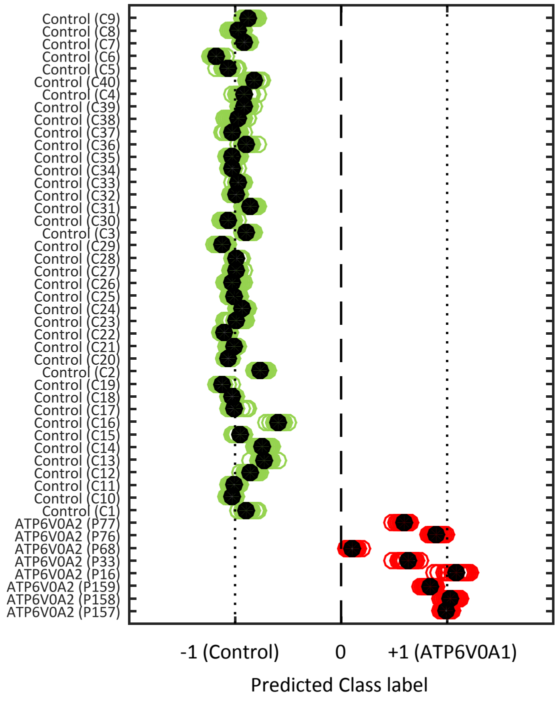

### Supplementary figure 4 - continued

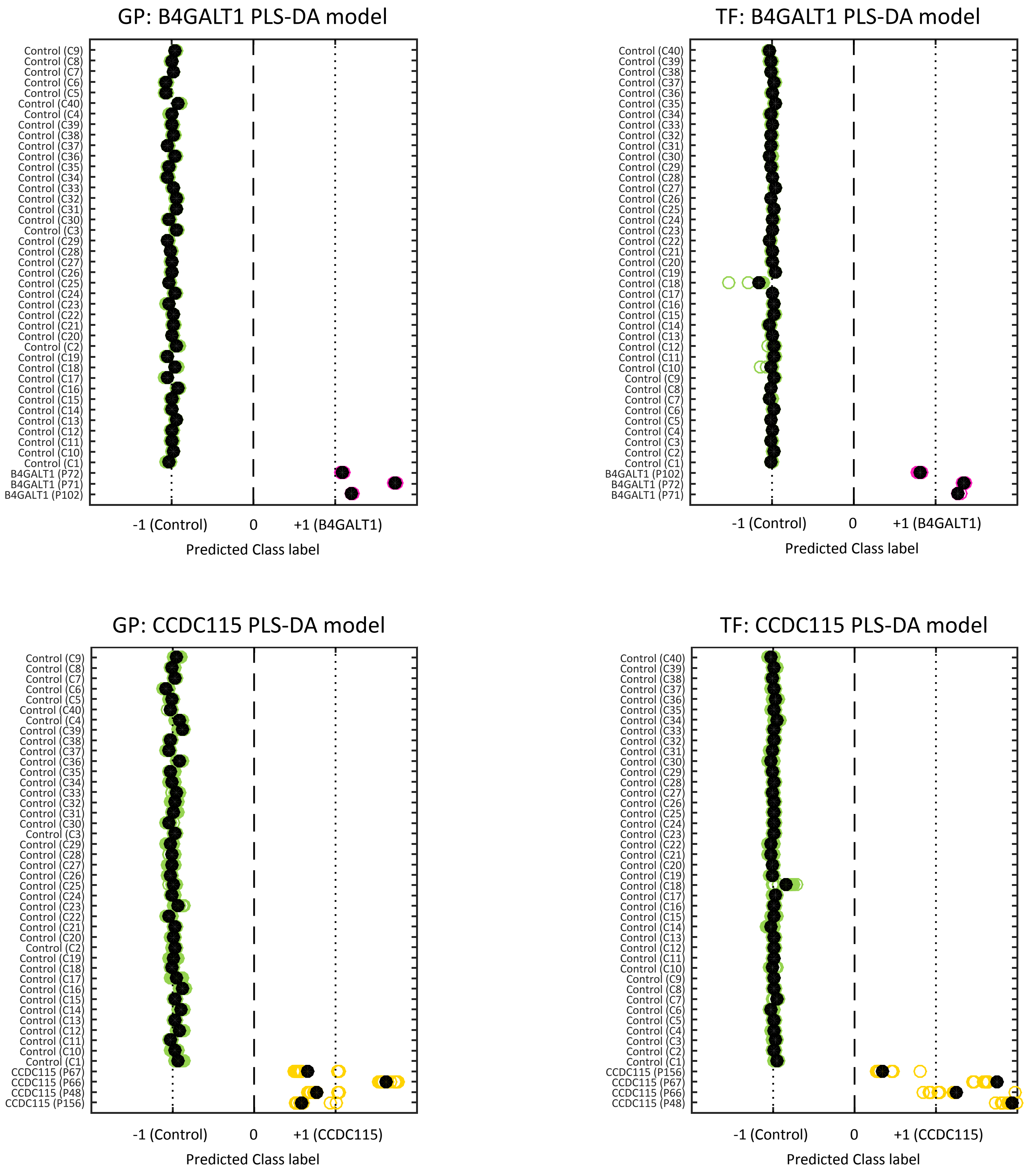

### Supplementary figure 4 - continued

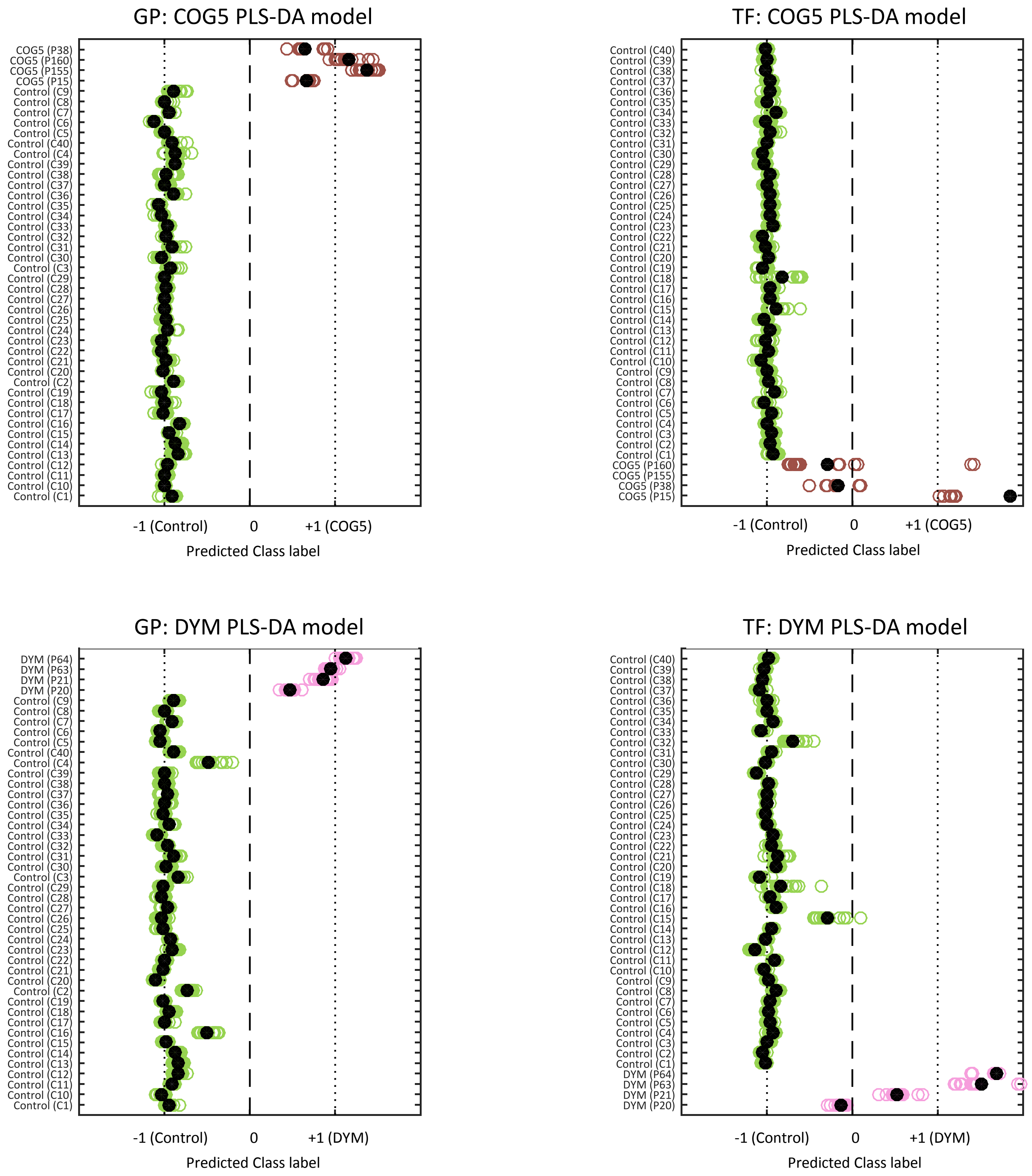

### Supplementary figure 4 - continued

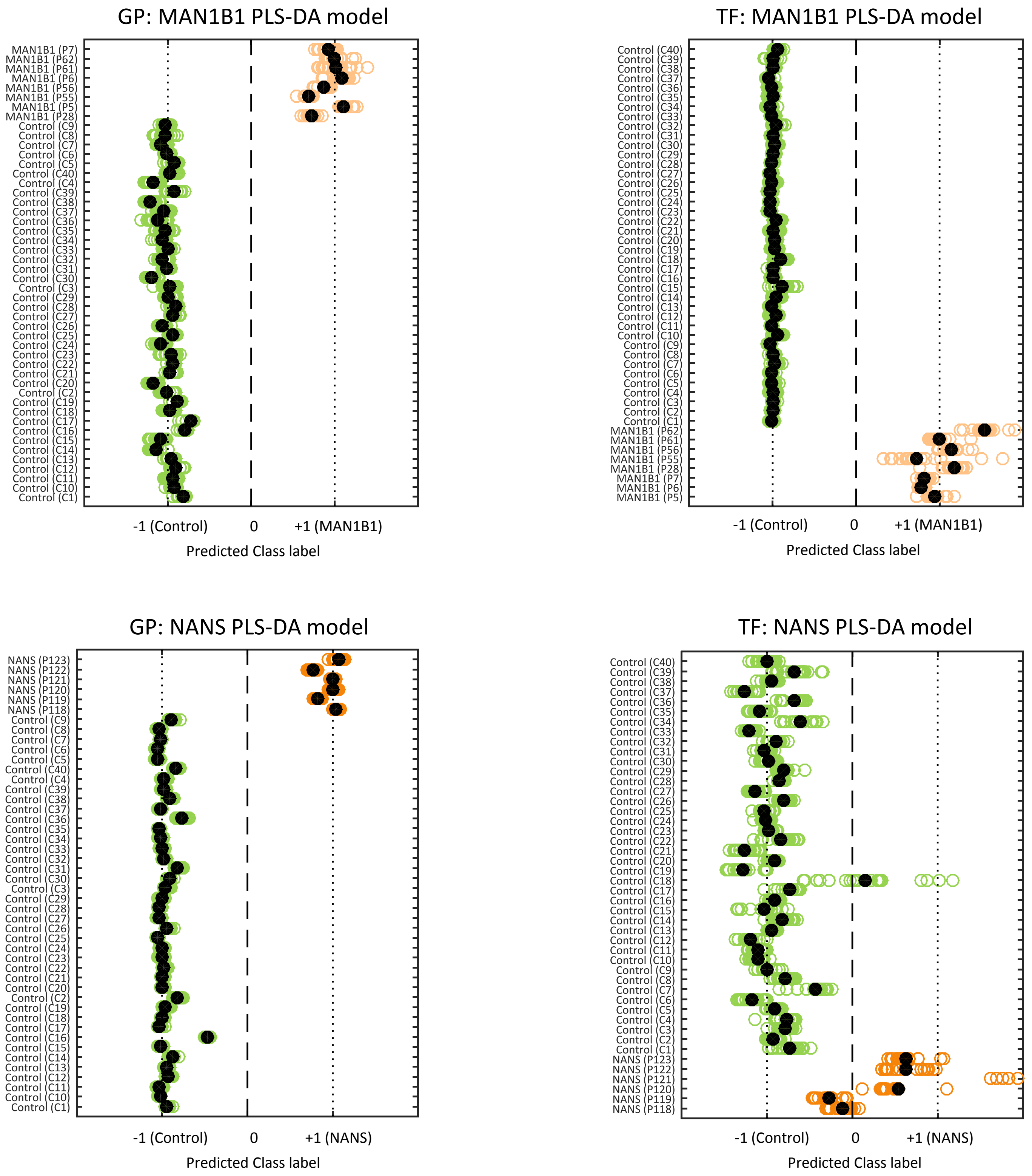

### Supplementary figure 4 - continued

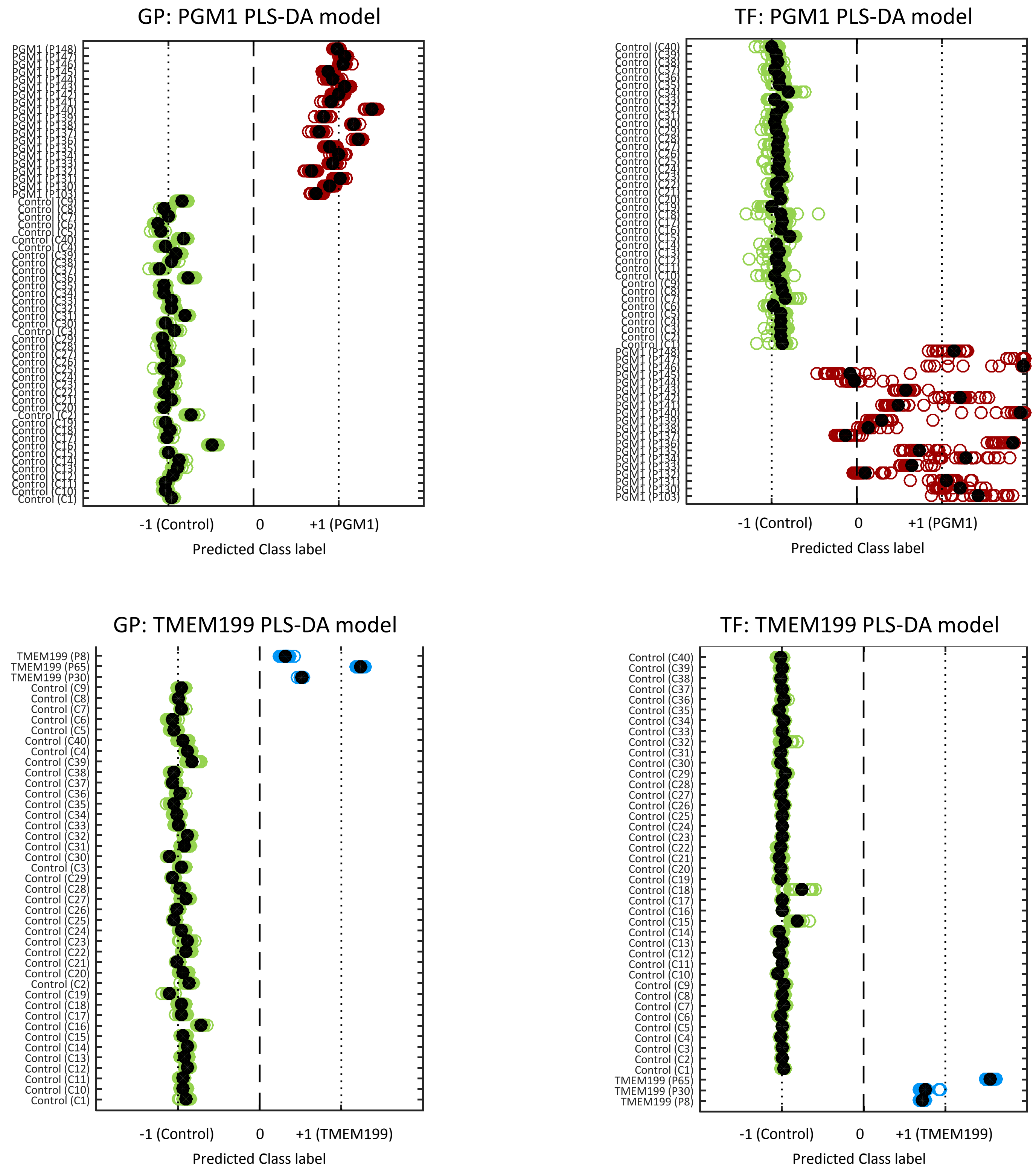

Supplementary figure 5a – differential chord diagrams B4GALT1

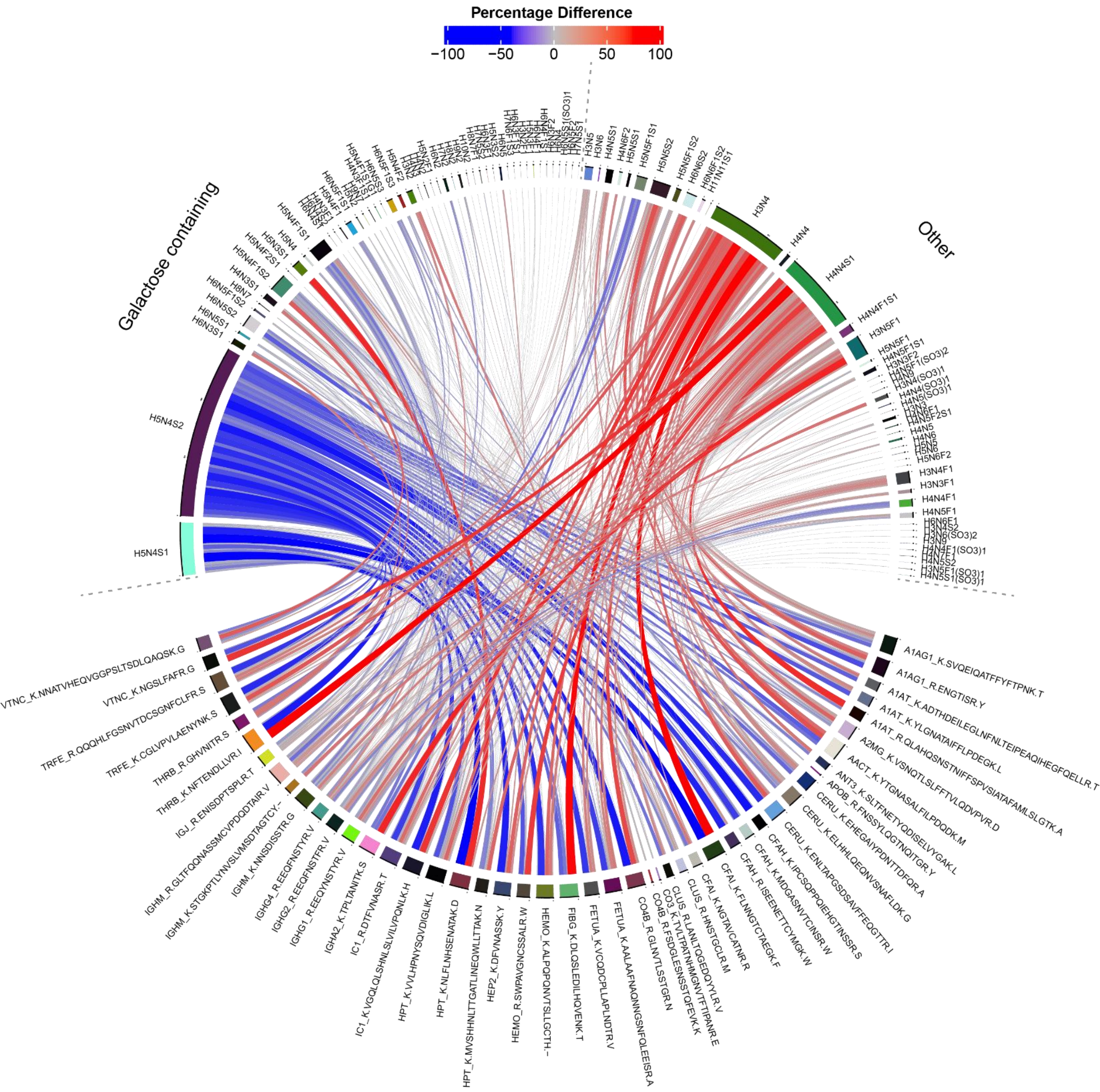


**Supplementary figure 5a.** Differential chord diagram to visualize glycoform (top) and glycan class (bottom) changes in the glycoproteome of B4GALT1 patients (n=3) versus healthy controls (n=40).

Supplementary figure 5b – microheterogeneity and macroheterogeneity data B4GALT1

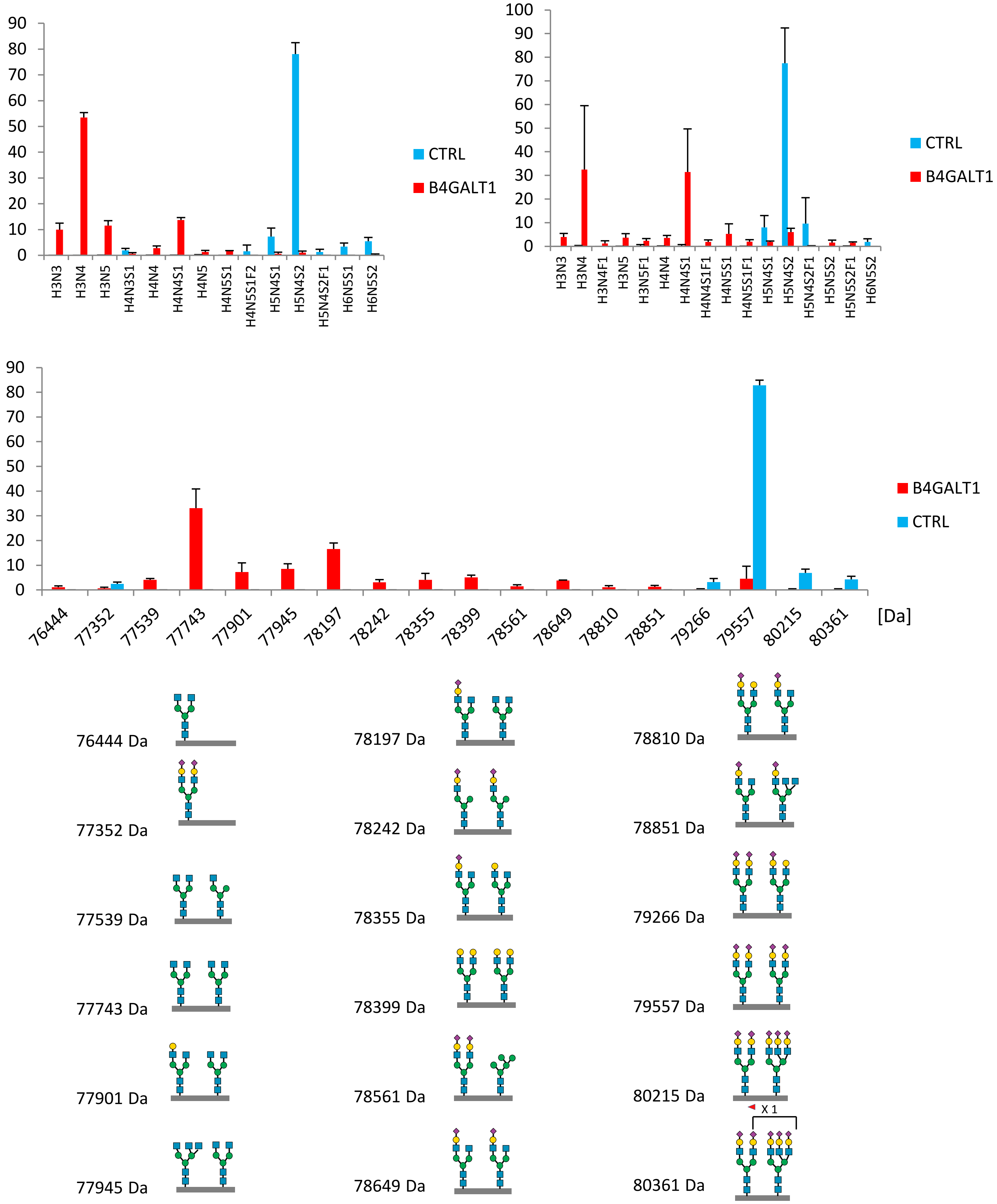


**Supplementary figure 5b.** Microheterogeneity and macroheterogeneity profiles of transferrin in B4GALT1 CDG patients (n=3) and healthy controls (n=40). Top: Microheterogeneity profiles of transferrin at Asn432 and Asn630 determined by glycopeptide profiling. Bottom: Macroheterogeneity profiles of transferrin determined by intact transferrin LC-MS. Intact transferrin masses with corresponding glycan sequences are provided to indicate likely combinations of glycans and do not represent site-specific annotations.

Supplementary figure 5c – differential chord diagram MAN1B1

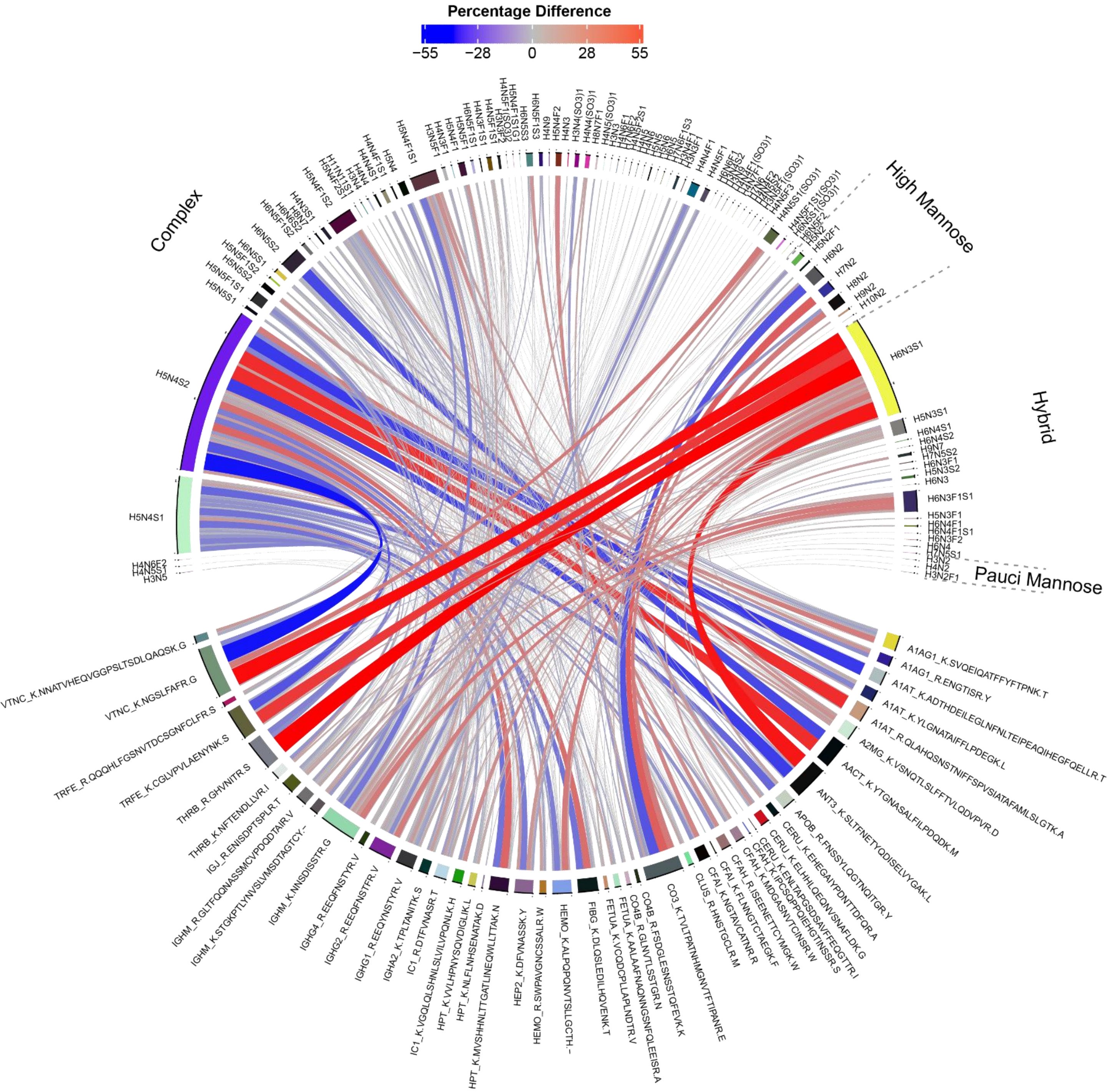


**Supplementary figure 5c.** Differential chord diagram to visualize glycoform changes in the glycoproteome of MAN1B1 patients (n=8) versus healthy controls (n=40).

Supplementary figure 5d – microheterogeneity and macroheterogeneity data MAN1B1

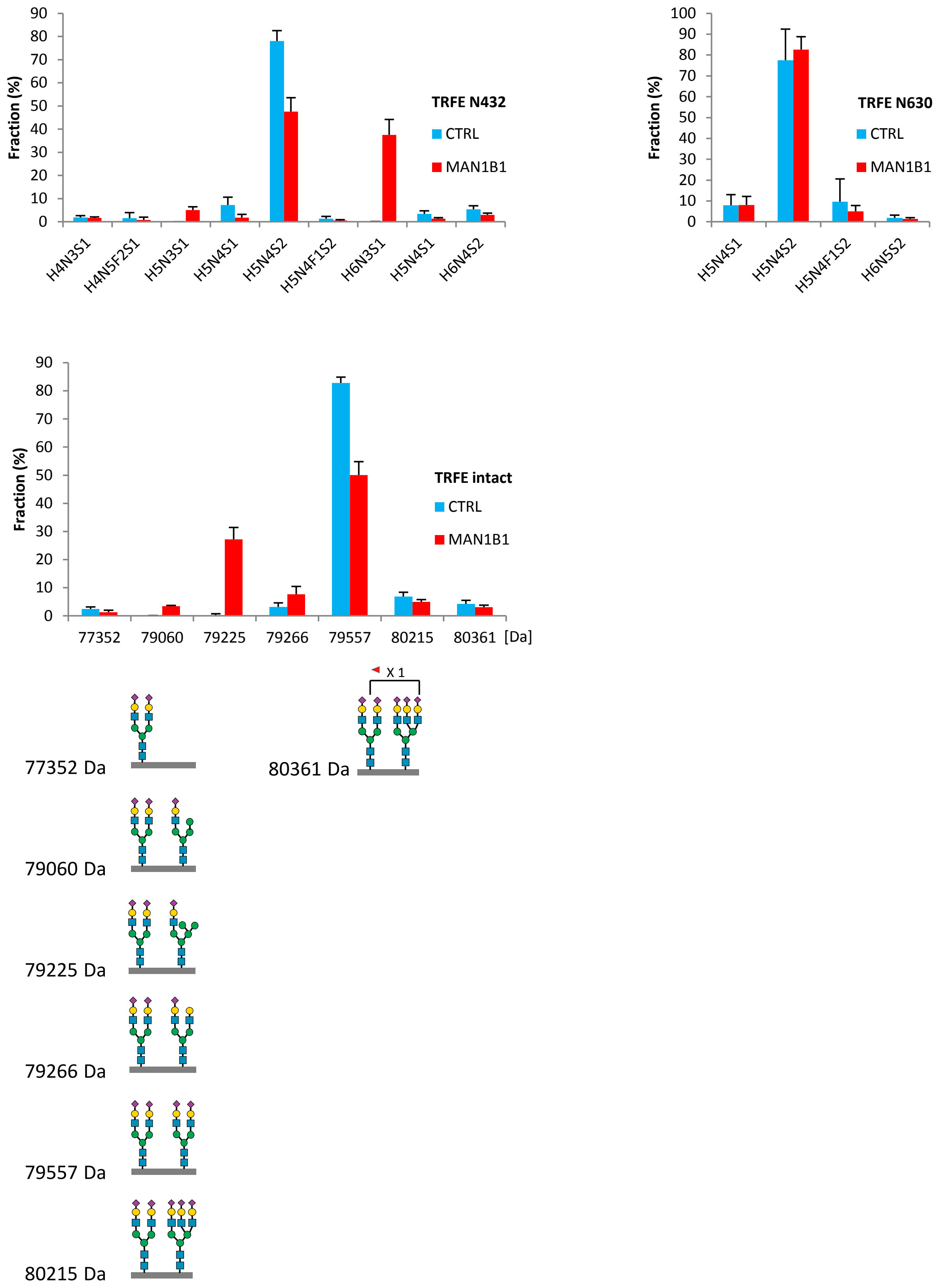


**Supplementary figure 5d.** Microheterogeneity and macroheterogeneity profiles of transferrin in MAN1B1 CDG patients (n=3) and healthy controls (n=40). Top: Microheterogeneity profiles of transferrin at Asn432 and Asn630 determined by glycopeptide profiling. Bottom: Macroheterogeneity profiles of transferrin determined by intact transferrin LC-MS. Intact transferrin masses with corresponding glycan sequences are provided to indicate likely combinations of glycans and do not represent site-specific annotations.

Supplementary figure 5e – differential chord diagrams SLC35A1

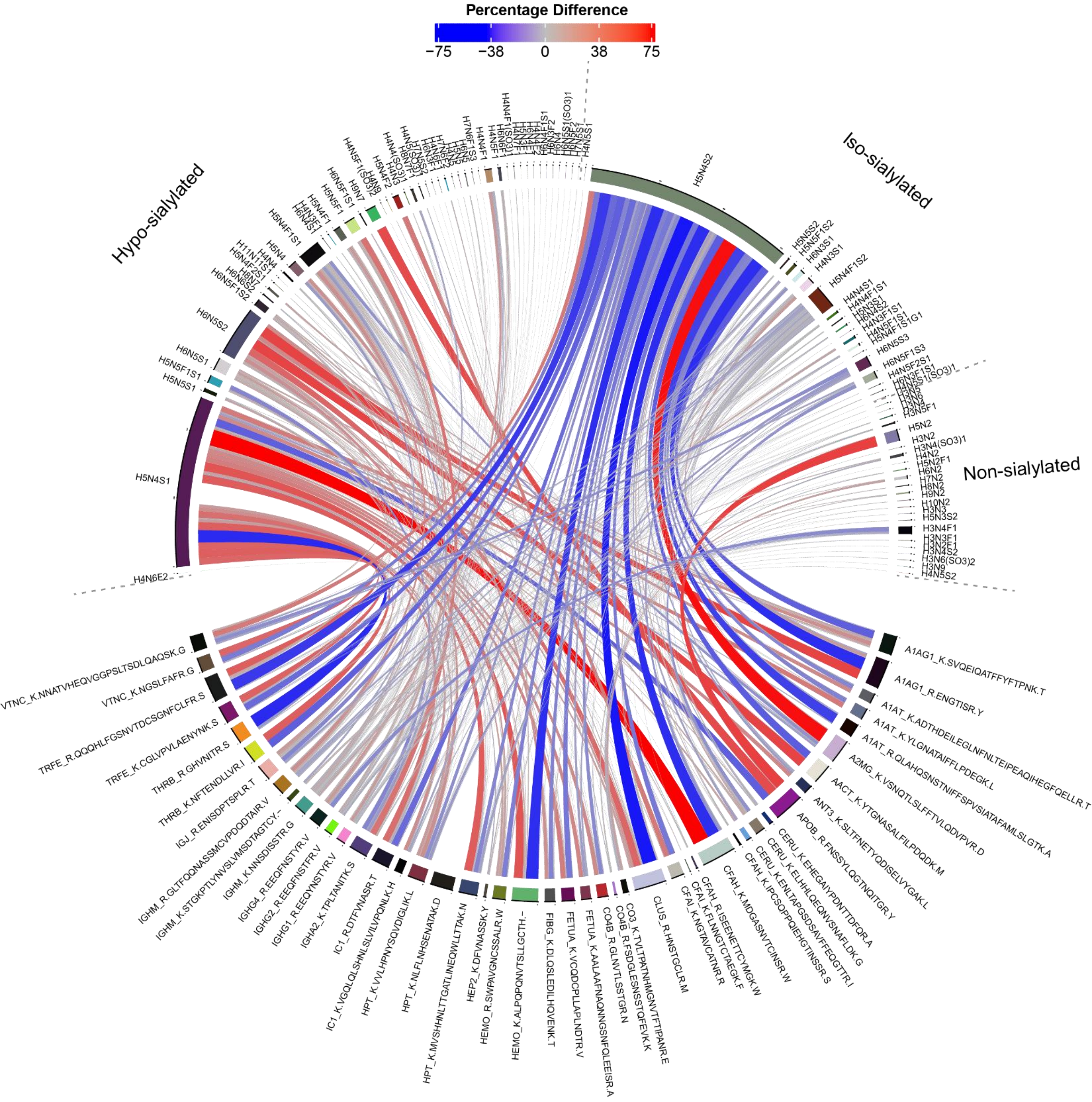


**Supplementary figure 5e.** Differential chord diagram to visualize glycoform (top) and glycan class (bottom) changes in the glycoproteome of an SLC35A1 patient (n=1) versus healthy controls (n=40).

Supplementary figure 5f – microheterogeneity and macroheterogeneity data SLC35A1

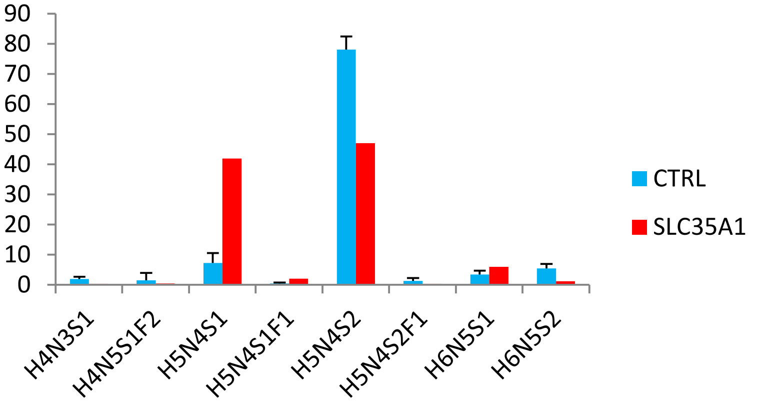


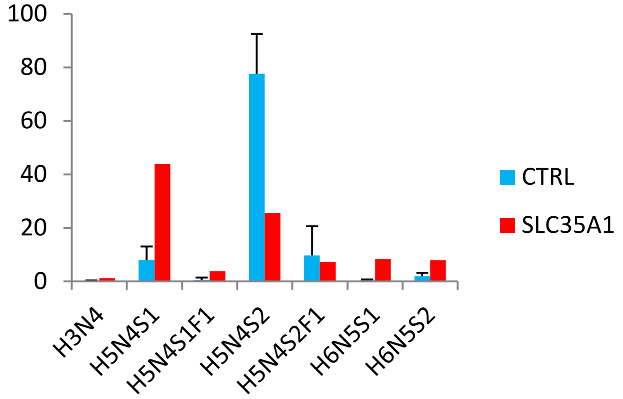


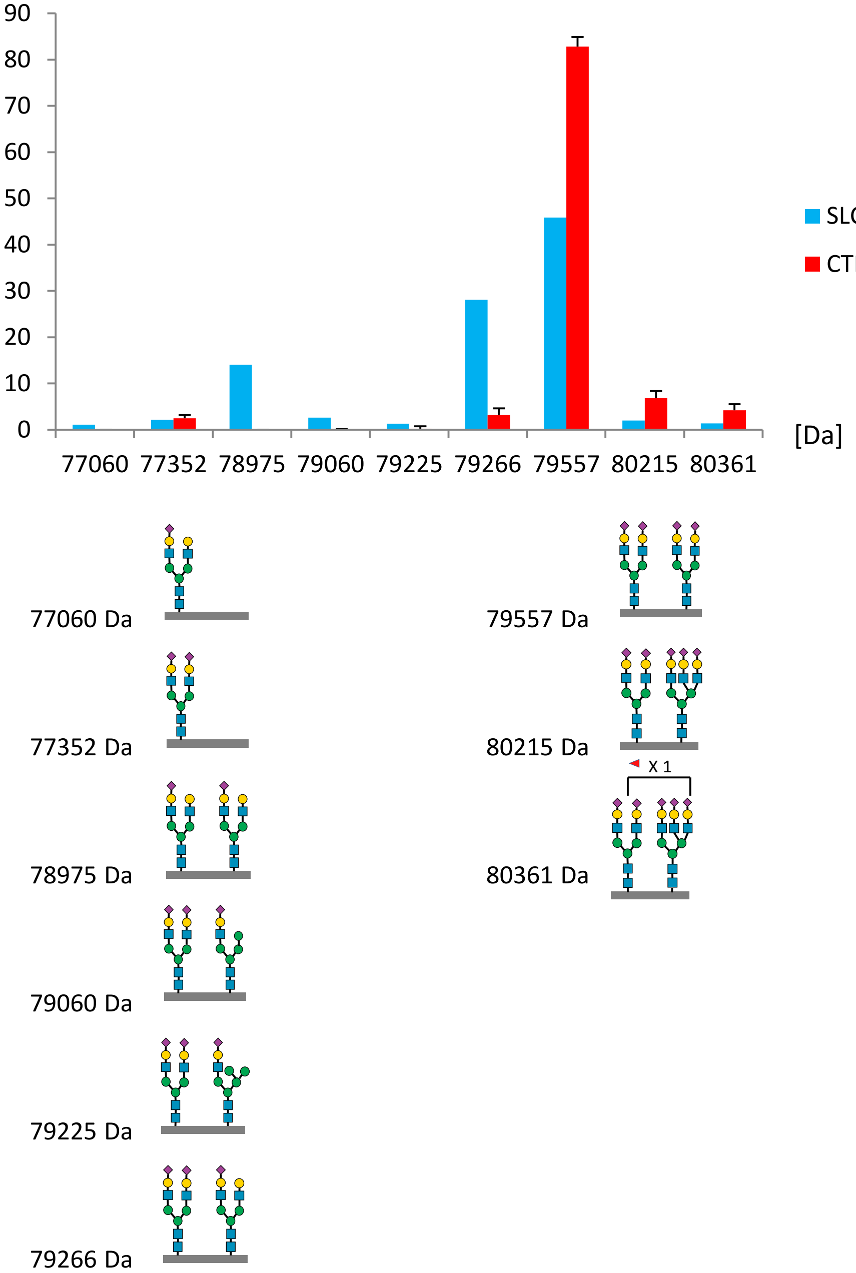


**Supplementary figure 5f.** Microheterogeneity and macroheterogeneity profiles of transferrin in a SLC35A1 CDG patient (n=1) and healthy controls (n=40). Top: Microheterogeneity profiles of transferrin at Asn432 and Asn630 determined by glycopeptide profiling. Bottom: Macroheterogeneity profiles of transferrin determined by intact transferrin LC-MS. Intact transferrin masses with corresponding glycan sequences are provided to indicate likely combinations of glycans and do not represent site-specific annotations.

Supplementary figure 5g – differential chord diagrams SLC35A3

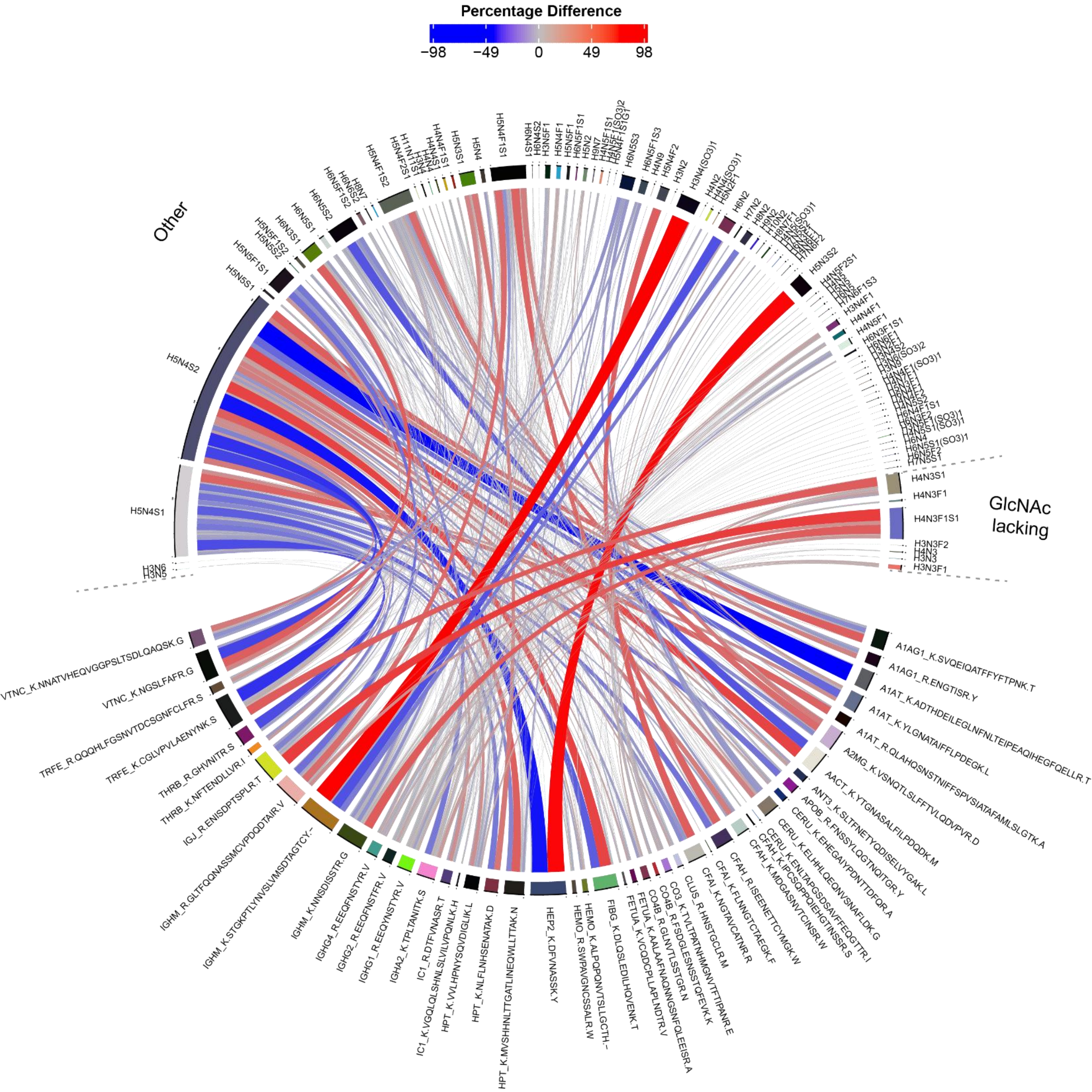


**Supplementary figure 5g.** Differential chord diagram to visualize relative glycoform (top) and glycan class (bottom) changes in the glycoproteome of an SLC35A3 patient (n=1) versus healthy controls (n=40).

Supplementary figure 5h – microheterogeneity and macroheterogeneity data SLC35A3

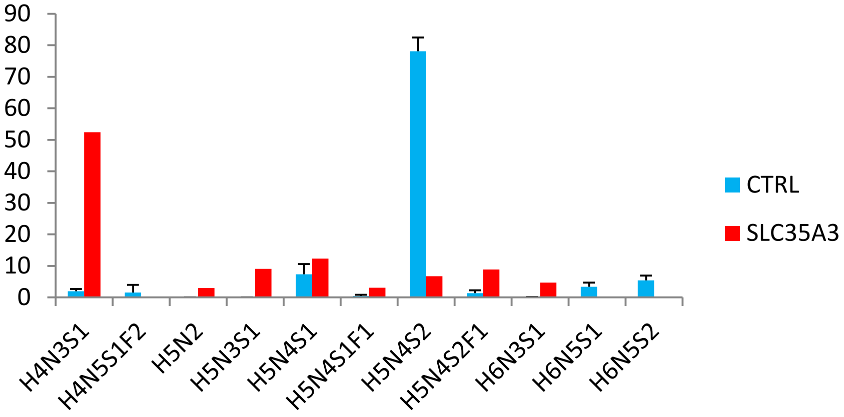


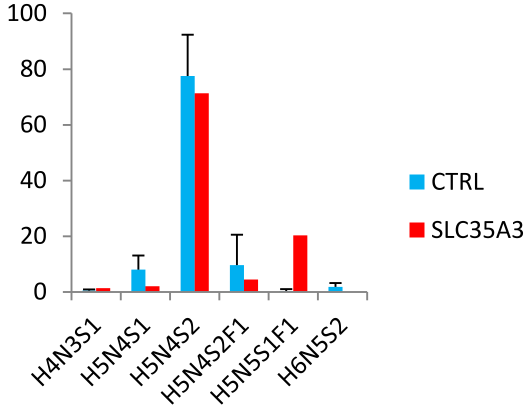


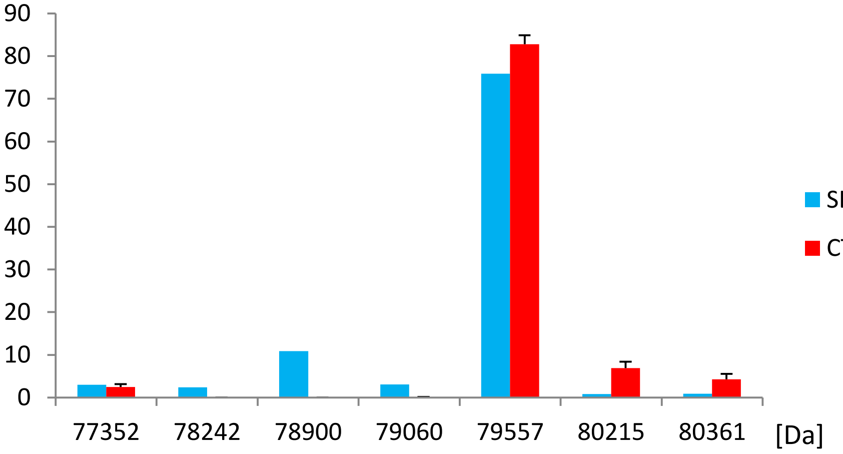


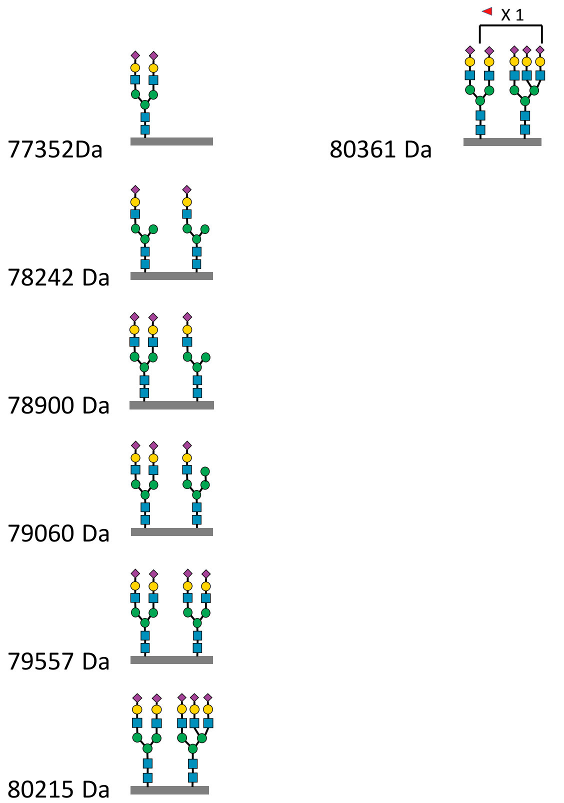


**Supplementary figure 5h.** Microheterogeneity and macroheterogeneity profiles of transferrin in a SLC35A3 CDG patient (n=1) and healthy controls (n=40). Top: Microheterogeneity profiles of transferrin at Asn432 and Asn630 determined by glycopeptide profiling. Bottom: Macroheterogeneity profiles of transferrin determined by intact transferrin LC-MS. Intact transferrin masses with corresponding glycan sequences are provided to indicate likely combinations of glycans and do not represent site-specific annotations.

Supplementary figure 5i – differential chord diagram SLC35C1

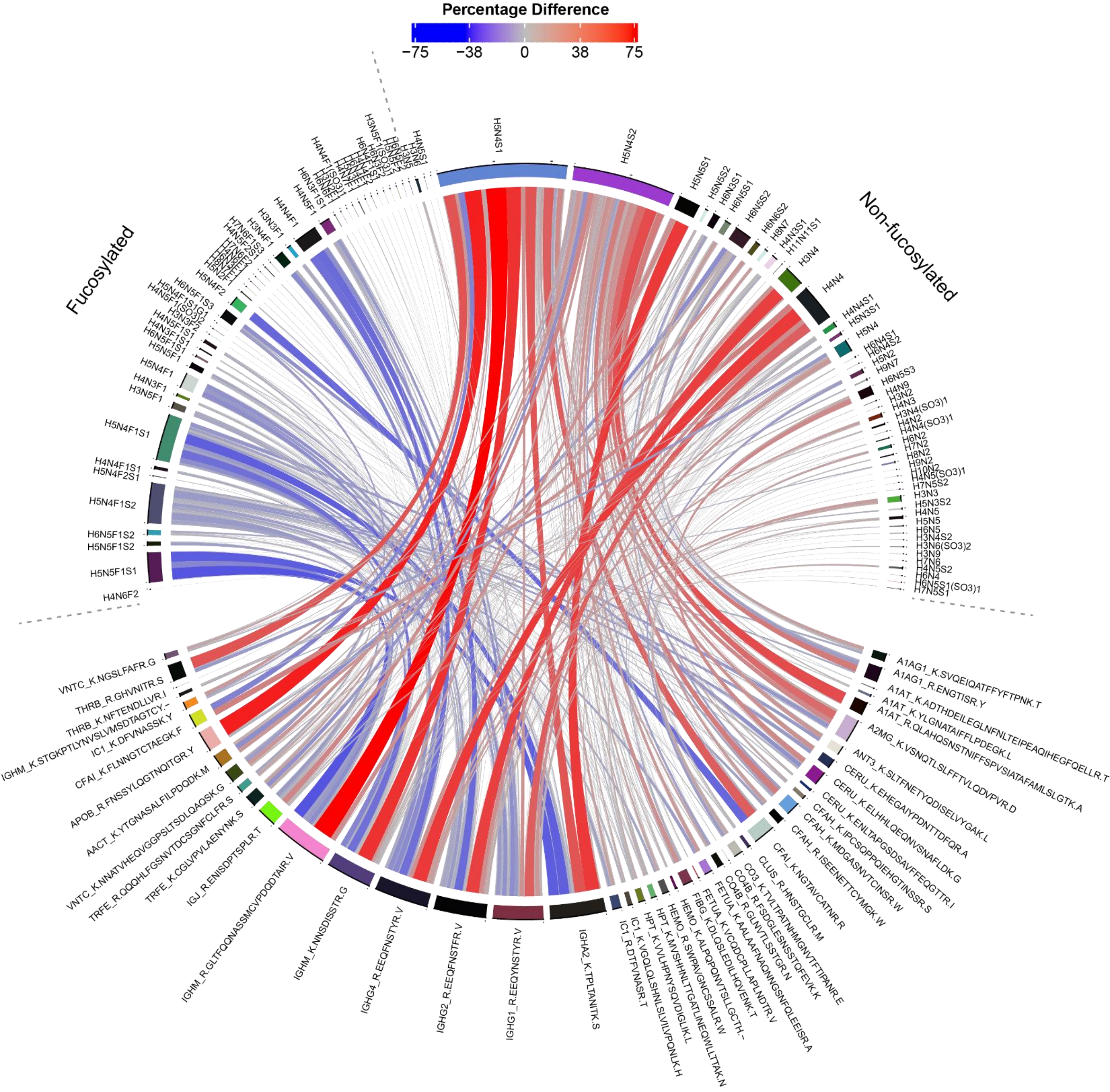


**Supplementary figure 5i.** Differential chord diagram to visualize relative glycoform changes in the glycoproteome of an SLC35C1 patient (n=1) versus healthy controls (n=40).

Supplementary figure 5j – microheterogeneity and macroheterogeneity data SLC35C1

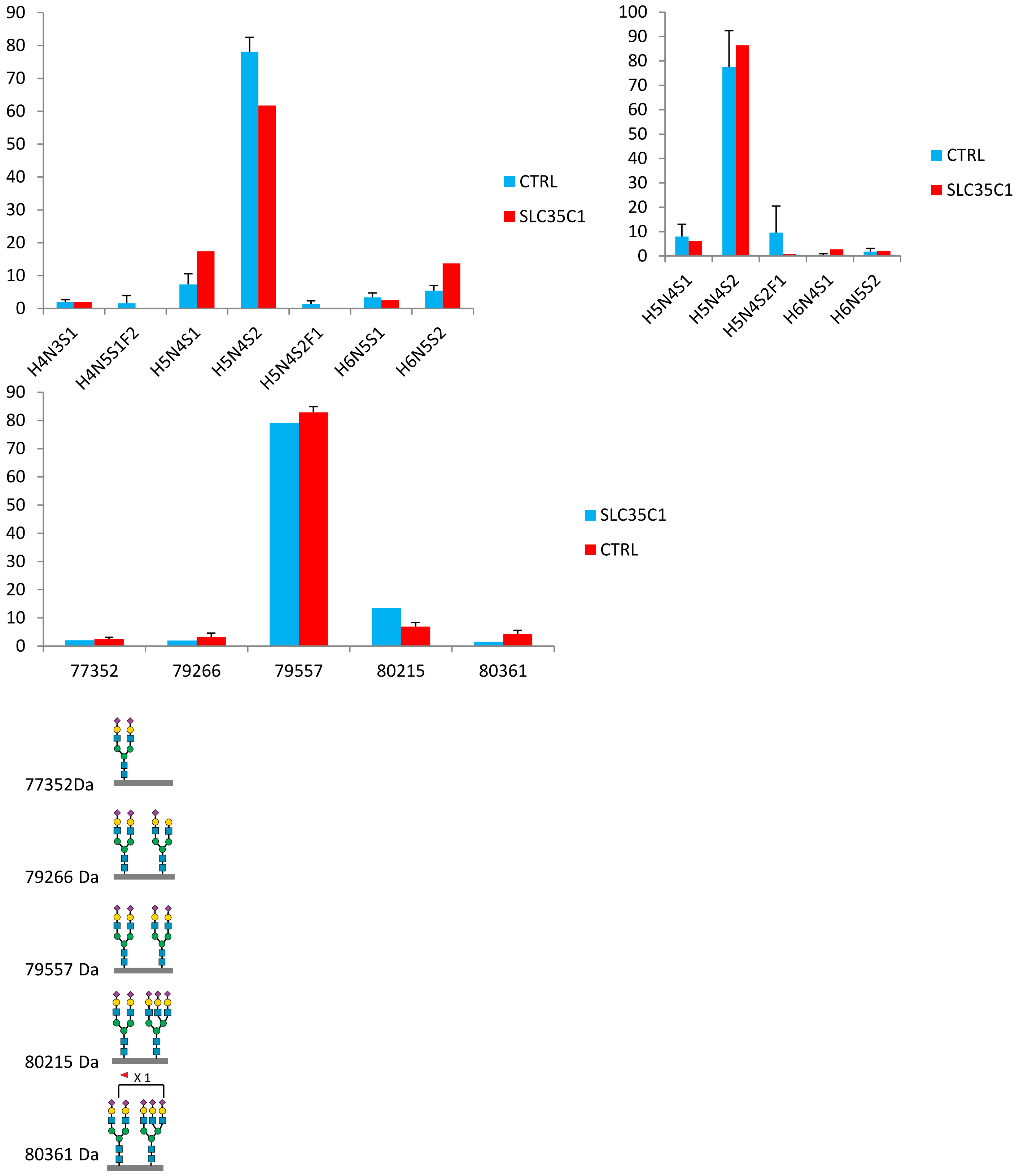


**Supplementary figure 5j.** Microheterogeneity and macroheterogeneity profiles of transferrin in a SLC35C1 CDG patient (n=1) and healthy controls (n=40). Top: Microheterogeneity profiles of transferrin at Asn432 and Asn630 determined by glycopeptide profiling. Bottom: Macroheterogeneity profiles of transferrin determined by intact transferrin LC-MS. Intact transferrin masses with corresponding glycan sequences are provided to indicate likely combinations of glycans and do not represent site-specific annotations.
